# The correct temporal connectivity of the DG-CA3 circuits involved in declarative memory processes depends on Vangl2-dependent planar cell polarity signaling

**DOI:** 10.1101/2024.05.28.596141

**Authors:** Noémie Depret, Marie Gleizes, Maïté Marie Moreau, Sonia Poirault-Chassac, Anne Quiedeville, Steve Dos Santos Carvalho, Vasika Venugopal, Alice Shaam Al Abed, Gael Barthet, Christophe Mulle, Aline Desmedt, Aline Marighetto, Claudia Racca, Mireille Montcouquiol, Nathalie Sans

## Abstract

In the hippocampus, dentate gyrus granule cells connect to CA3 pyramidal cells via their axons, the mossy fibers (Mf). The synaptic terminals of Mfs (Mf boutons, MfBs) form large and complex synapses with thorny excrescences (TE) on the proximal dendrites CA3 pyramidal cells (PCs). MfB/TE synapses have distinctive “detonator” properties due to low intitial release probability and large presynaptic facilitation. The molecular mechanisms shaping the morpho-functional properties of MfB/TE synapses are still poorly understood, though alterations in their morphology are associated with Down syndrome, intellectual disabilities, and Alzheimer’s disease. Here, we identify the core PCP gene *Vangl2* as essential to the morphogenesis and function of MfB/TE synapses. Vangl2 colocalises with the presynaptic heparan sulfate proteoglycan glypican 4 (GPC4) to stabilise the postsynaptic orphan receptor GPR158. Embryonic loss of Vangl2 disrupts the morphology of MfBs and TEs, impairs ultrastructural and molecular organisation, resulting in defective synaptic transmission and plasticity. In adult, the early loss of Vangl2 results in a number of hippocampus-dependent memory deficits including characteristic flexibility of declarative memory, organisation and retention of working/ everyday-like memory. These deficits also lead to abnormal generalisation of memories to salient cues and diminished ability to form detailed contextual memories. Together, these results establish Vangl2 as a key regulator of DG-CA3 connectivity and functions.

**Highlights:**

- Vangl2 is a key regulator of MfB/TE synapses morphogenesis and plasticity in CA3
- Vangl2-mediated GPC4-GPR158 interaction maintains MfB-TE pre-synaptic morphology and function
- *Vangl2* deletion affects declarative memory in adult mice
- Vangl2 function is necessary for contextual learning and its loss leads toa maladaptive fear memory for salient cues

## Introduction

Correct morpho-functional development of the circuit assembled by the axon (mossy fiber or Mf) boutons (MfB) of granule cells (GC) of the dentate gyrus (DG) and the postsynaptic dendritic thorny excrescences (TE) of CA3 pyramidal cells (PC) in the *stratum lucidum* (*sl*) is critical for hippocampal function. Alterations in the morphology and/or function of MfB/TE synapses are associated with several central nervous system pathologies, including Down syndrome, intellectual disabilities ^1,2^, or Alzheimer’s disease ^3,4^. However, little is known about the molecular and cellular basis of this morphogenesis, sculpting these large MfB/TE synapses and contributing to their distinctive “detonator” properties ^5–7^. This morphogenesis takes place in the early post-natal stages ^5,6^, and is restricted spatially. In fact, the synapses on the medial portion of the same apical dendrites in the *stratum radiatum* (*sr*) are smaller, less complex, and with milder facilitation.

Recent studies have identified the neurexin/neuroligin complex, the leucine-rich repeat transmembrane (LRRTM) family, and the heparan sulphate proteoglycan (HSPG) cell surface molecules as critical for MfB/TE development and function ^8–10^. Glypican 4 (GPC4) and its interactor GPR158 were notably identified as critical for the proper development of functional MfB/TE synapses ^10^. Changes in GPC4 levels play a role in the sprouting of MfB/TE synapses in the adult, in a mouse model of temporal lobe epilepsy ^11^. Adhesion molecules and the cytoskeleton also participate in the growth and morphogenesis of MfB/TE synapses ^8,12^. A recent study from our group showed that Vangl2 protein, a Wnt/PCP molecule, is enriched in specific regions of the adult mouse hippocampus, and that the postnatal deletion of the *Vangl2* gene delays maturation of GCs and disrupts specific memory processes ^13^. The Wnt/PCP pathway is a conserved signaling pathway that shapes tissues during development by regulating adhesion complexes and the cytoskeleton. In mammals, including humans, mutations in *VANGL2* (OMIM 600533) lead to severe morphological deficits of the nervous system known as neural tube defects. We and others have shown that *Vangl2* is involved in several early and late mechanisms of brain maturation and function ^14,15^. Here we show that Vangl2 is enriched in the *sl* during early postnatal stages and modulates the early growth and morphogenesis of MfB/TE synapses. We propose that Vangl2 participates in the GPC4/GPR158 pathway that controls MfB/TE morphogenesis and function. In support of our hypothesis, we show that early deletion of *Vangl2* results in the disruption of the structural development and plasticity of MfB/TE synapses, as well as of presynaptic physiological properties. In adult mice, these deficits lead to the impairment of several hippocampus-dependent cognitive processes, including memory flexibility, organisation and retention of working/everyday-like memory and contextual fear memory.

## Results

### Vangl2 spatiotemporal profile correlates with MfB/TE synapse morphogenesis

Mfs first contact CA3 pyramidal cells around 7 days after birth (P7) ^7^. At this stage, Vangl2 is strongly expressed throughout the hippocampus, in particular in the DG and in the developing Mfs that originate from the GCs of the DG (Figure 1A, B). Between P7 and P14, when the morphogenesis of the MfB/TE giant synapse takes place, Vangl2 accumulates in the *sl* of the CA3 (Figure 1B). The *sl* is a region where the Mfs contact the proximal apical dendrites of CA3 PCs. This maturation continues until P21 (Figure 1B), when MfB/TE synapses are considered to be structurally mature ^7,16^. We generated a conditional *Vangl2* knockout mouse (hereafter referred to as Emx1-Vangl2), and confirmed the absence of the protein in the hippocampus of Emx1-Vangl2 mice (Figure S1A, B) ^15^.

**Figure 1:**
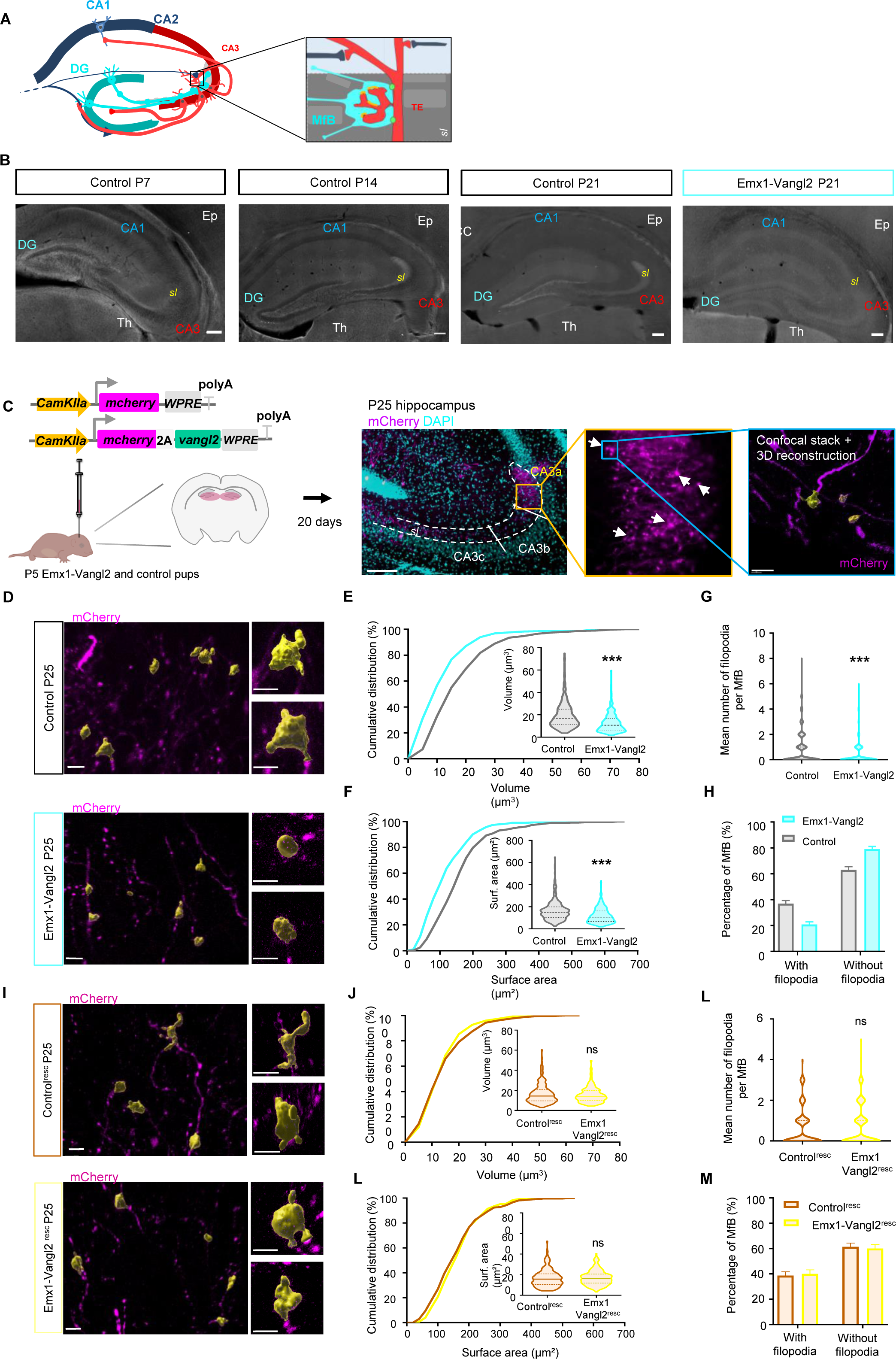
Vangl2 protein is enriched in the stratum lucidum and is necessary for the morphologenesis of presynaptic MfB. (**A**) Schematic representation of the hippocampal trisynaptic circuit between the dentate gyrus (DG), CA3, and CA1 with the schematic of the mossy fibre synapse (MfB/TE synapse) formed by the presynaptic mossy fibre bouton (MfB) of the granule cells and the postsynaptic thorny excrescence (TE) of the pyramidal neurons in the *stratum lucidum* (sl) of the CA3. **(B)** Immunolabelling for endogenous Vangl2 in mouse brains shows that Vangl2 is expressed across the entire hippocampus, with a specific enrichment in the granular and subgranular layer of the DG and the mossy fibre projections of the granule cells at P7. This expression pattern is still present in P14 mouse brain, with strong enrichment in the sl of the CA3. At P21, Vangl2 protein expression is more confined than in P14 mice brains but Vangl2 is enriched in the DG subgranular zone and the sl. Note that Vangl2 is also expressed in the ependymal cells lining the ventricles (Ep), as well as in more ventral regions such as the thalamus (Th). In the mutant, Vangl2 labeling is absent in the hippocampus but can still be seen in the thalamus and the ependymal cells. Scale bar, 200 µm. **(C)** Experimental design for stereotaxic virus injections in the DG of P5 pups and representative confocal image of labelled presynaptic MfB in the sl of P25 control mice in magenta, with the subsequent 3D reconstruction of MfB in yellow. Scale bar, 200µm and 5µm. **(D,I)** Representative confocal images of labelled MfB (magenta) with 3D reconstruction (yellow) in the sl of P25 Emx1-Vangl2 and control mice (D) or of P25 control^resc^ and Emx1-Vangl2^resc^ mice (I). Scale bar, 5 µm. **(E-H)** Quantifications of volume (E), surface area (F), number of filopodia per MfB (G), and percentage of MfB with filopodia (H) in P25 Emx1-Vangl2 and control mice. N control= 271 MfB from 7 mice, N Emx1-Vangl2 = 293 MfB from 7 mice. Mann-Whitney test, ***p < 0.001. **(J-M)** Quantifications of volume (J), surface area (K), number of filopodia per MfB (L), and percentage of MfB with filopodia (M) in P25 Emx1-Vangl2^resc^ and control^resc^ mice. N control^resc^ = 263 MfB from 8 mice, N Emx1-Vangl2^resc^ = 245 MfB from 8 mice. Mann-Whitney test, ns p>0.05. Error bars represent SEM. From E to H and from J to M data are presented as median with 25th and 75th percentile. The shaded area represents the probability distribution of the variable.

Vangl2 colocalises with doublecortin (DCX), which labels Mf projections arising from immature GCs, as well as with Tau (an axonal marker), however it appears to not colocalise with ZnT3 (a marker of MfBs), and PSD-95 (a marker of postsynaptic densities) (Figure S1C). These data show that the expression profile of Vangl2 correlates with the timing of morphological maturation of MfB/TE giant synapses.

### Vangl2 participates in the morphogenesis of the presynaptic compartments of MfB/TE giant synapse

We infected the DG GCs of postnatal day 5 (P5) Emx1-Vangl2 mice and of control littermates that did not express the Cre recombinase with an AAV2/9-CaMKII(0.4)-mCherry-WPRE virus in order to achieve sparse labelling of neurons (Figure 1C). We quantified the morphological parameters of the 3D reconstructed MfBs located in the *sl* of P25 mice (Figure 1C). We observed a significant decrease in the volume of the MfB (Figure 1E; control: 18.28 ± 0.74 µm^3^, Emx1-Vangl2: 13.79±0.50 µm^3^; Mann-Whitney test ***p<0.0001) and surface area (Figure 1F; control: 171.17±5.70 µm², Emx1-Vangl2: 135.66±4.23 µm²; Mann-Whitney test ***p<0.0001) in Emx1-Vangl2 mice compared to controls. Emx1-Vangl2 mice had fewer filopodia per MfB than controls (Figure 1G; control: 0.82±0.08 filopodia, Emx1-Vangl2: 0.36±0.05 filopodia; Mann-Whitney test ***p<0.0001). Furthermore, we observed a significant reduction in the percentage of filopodia-bearing MfBs in P25 Emx1-Vangl2 mice compared to controls (Figure 1I; control: 38.28±2.96%, Emx1-Vangl2: 22.87±2.46%; Mann-Whitney test ***p<0.0001). The data show that in absence of Vangl2, MfBs are structurally smaller, less convoluted, and with fewer filopodia: all characteristic of immature MfBs. The integrated fluorescent signals for synaptoporin, bassoon, and synapsin1 (presynaptic markers) were significantly reduced in *sl* mutants compared to controls (Figure S2A-G; see methods) (synaptoporin: −18.11±2.57%, One sample t-test ***p=0.0001; bassoon: −10.11±1.56%, One sample t-test ***p=0.0002; synapsin1: −16.70±3.23%, One sample t-test **p= 0.0021). These results support the notion that MfBs are immature in the absence of Vangl2.

The early injection of an AAV virus allowing the expression of Vangl2 in DG GCs in P5 Emx1-Vangl2 mice and controls (control^resc^ and Emx1-Vangl2^resc^) (Figure 1C) fully rescued the morphogenesis of MfB/TE synapses, confirming the necessity for Vangl2 expression. At P25 we observed no difference between the MfBs of Emx1-Vangl2^resc^ and control^resc^ mice (Figure 1I). There was no statistical difference in volume (Figure 1J; control^resc^: 16.43±0.59 µm^3^, Emx1-Vangl2^resc^: 15.60±0.50 µm^3^; Mann-Whitney test ns p= 0. 76), nor in the surface area (Figure 1K; control^resc^: 164.75±5.09 µm^2^, Emx1-Vangl2^resc^: 169.96±4.51µm^2^; Mann-Whitney test ns p= 0.17) between the MfBs of control^resc^ and Emx1-Vangl2^resc^. Furthermore, the number of filopodia per MfB and the percentage of MfBs with filopodia were also rescued by Vangl2 re-expression (Figure 1L: control^resc^: 0.62±0.058 filopodia, Emx1-Vangl2^resc^: 0.67±0.06 filopodia; Mann-Whitney test ns p=0.60; Figure 1M: control^resc^: 38.64±3.00%, Emx1-Vangl2^resc^: 40±3.14%; Mann-Whitney test ns p=0.75). Of note, the morphological parameters of control^resc^ mice were not different between genotypes, demonstrating that re-expression of exogenous Vangl2 was not disruptive at 3 weeks (Figure S3A-D). These results show that re-expression of Vangl2 in the GCs rescues the morphological deficits induced by early *Vangl2* deletion. Altogether, the data demonstrated that Vangl2 is necessary for the morphological and molecular maturation of MfB/TE synapses.

### Vangl2 participates in the morphogenesis of the pre- and postsynaptic compartments of MfB/TE giant synapse

To evaluate the pre- and postsynaptic compartment at the nanoscale and in 3D, we used serial block face scanning electron microscopy (SBFSEM) in the *sl* of the CA3a region (Figure 2 and movie S1). We observed a reduction in the volume of TEs in Emx1-Vangl2 mice (control: 1.38±0.15 µm^3^, Emx1-Vangl2: 0.45±0.06 µm^3^; Mann-Whitney test ***p<0.0001), with a higher number of small TEs (< 0.1 µm^3^) in Emx1-Vangl2 mice compared to controls (Figure 2A and 2C). The presynaptic MfB volume was significantly reduced in Emx1-Vangl2 mice (Figure 2B and 2D) (control: 7.00±0.57 µm^3^, Emx1-Vangl2: 2.84±0.41 µm^3^; Mann-Whitney test ***p<0.0001). The number of postsynaptic densities (PSDs) per TE was significantly reduced in Emx1-Vangl2 mice, averaging 0.0056±0.36 PSDs in controls compared to 0.00228±0.19 PSDs in mutants (Figure 2E and 2F; Mann-Whitney test ***p<0.0001). The average volume of the PSD was stable between the genotypes (Figure 2E and 2G; control: 0.0057±0.001 µm^3^, Emx1-Vangl2: 0.0056±0.001 µm^3^; Mann-Whitney test ns p=0.7118), whereas the average surface area of the PSD was reduced in Emx1-Vangl2 mice (Figure 2E and 2H; control: 0.29±0.011 µm^2^, Emx1-Vangl2: 0.25±0.01 µm^2^; Mann-Whitney test **p = 0.0029). These results confirm and extend our previous results on the reduced size and complexity of the presynaptic compartment and show a parallel morphogenetic defect in the postsynaptic compartment of MfB/TE synapses in absence of Vangl2.

**Fig. 2:**
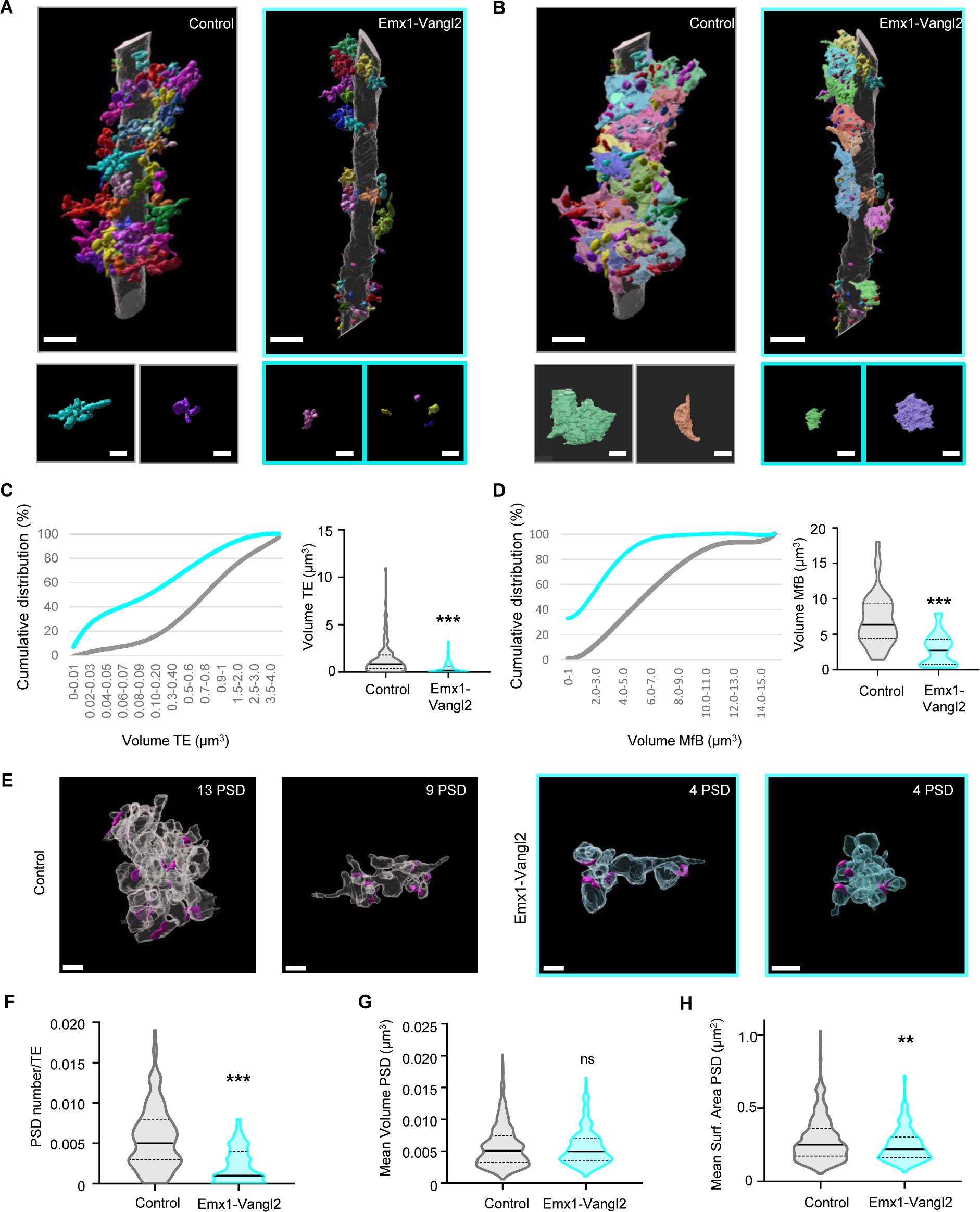
Size and complexity of MfB/TE synapses are reduced in Emx1-Vangl2 mice. **(A-B)** Representative 3D rendering of dendrites segmented from SBFSEM stacks of CA3a in P21 control and Emx1-Vangl2 animals. Scale bar, 3 µm. TE are represented in bold colors (A), MfBs are represented in pastel colors (B). Scale bar, 1 µm. **(C-D)** Cumulative distributions and quantifications of 3D reconstructed TEs (C) and MfBs (D) average volumes in P21 control (grey) and Emx1-Vangl2 (turquoise) mice. Number of TEs: Control: n = 119, Emx1-Vangl2: n = 117; Number of MfBs: control: n = 42, Emx-Vangl2: n = 30. Mann-Whitney test, ***p<0.001. **(E)** Representative 3D rendering of PSDs (pink) onto TEs segmented from SBFSEM stacks in P21 control and Emx1-Vangl2 animals. Scale bar, 1 µm. **(F-H)** Quantifications of the mean number of PSD per TE (F), mean PSD volume (G) and surface area (H) for control (grey) and Emx1-Vangl2 P21 mice (turquoise). Number of PSDs: control: n = 681, Emx1-Vangl2: n = 255. Mann-Whitney test, **p<0.01 ***p<0.001. From C to H, data are presented as median with 25th and 75th percentile. The shaded area represents the probability distribution of the variable.

### Vangl2 colocalises with presynaptic GPC4 to stabilise postsynaptic GPR158

The HSPG glypican 4 (GPC4) is a good candidate to transduce Vangl2 signaling at MfB/TE synapses ^10,17,18^ (Figure 3A). In COS-7 cells, the two proteins colocalised in vesicles or clusters and at the plasma membrane (Figure 3B and 3D). To assess whether the two proteins traffic together, we used a GPC4 variant (GPC4-myc) that remains blocked in the endoplasmic reticulum (ER)^10^ (Figure 3B). When co-expressed with Vangl2, both proteins remain in the ER (Figure 3E). These results indicate that GPC4 can associate with Vangl2 early in the trafficking pathway and that both proteins are trafficked together to the plasma membrane. Vangl2 did not colocalise with the orphan receptor GPR158, a transsynaptic interactor of GPC4, except at the Golgi apparatus in COS-7 cells (Figure 3C and 3F). In DIV17 (Days In Vitro) neurons, Vangl2 and GPC4 partially colocalised at the growth cone of the axon (Figure 3G). These data support the hypothesis that GPC4 and Vangl2 are associated in presynaptic compartments during axonal development.

**Fig. 3:**
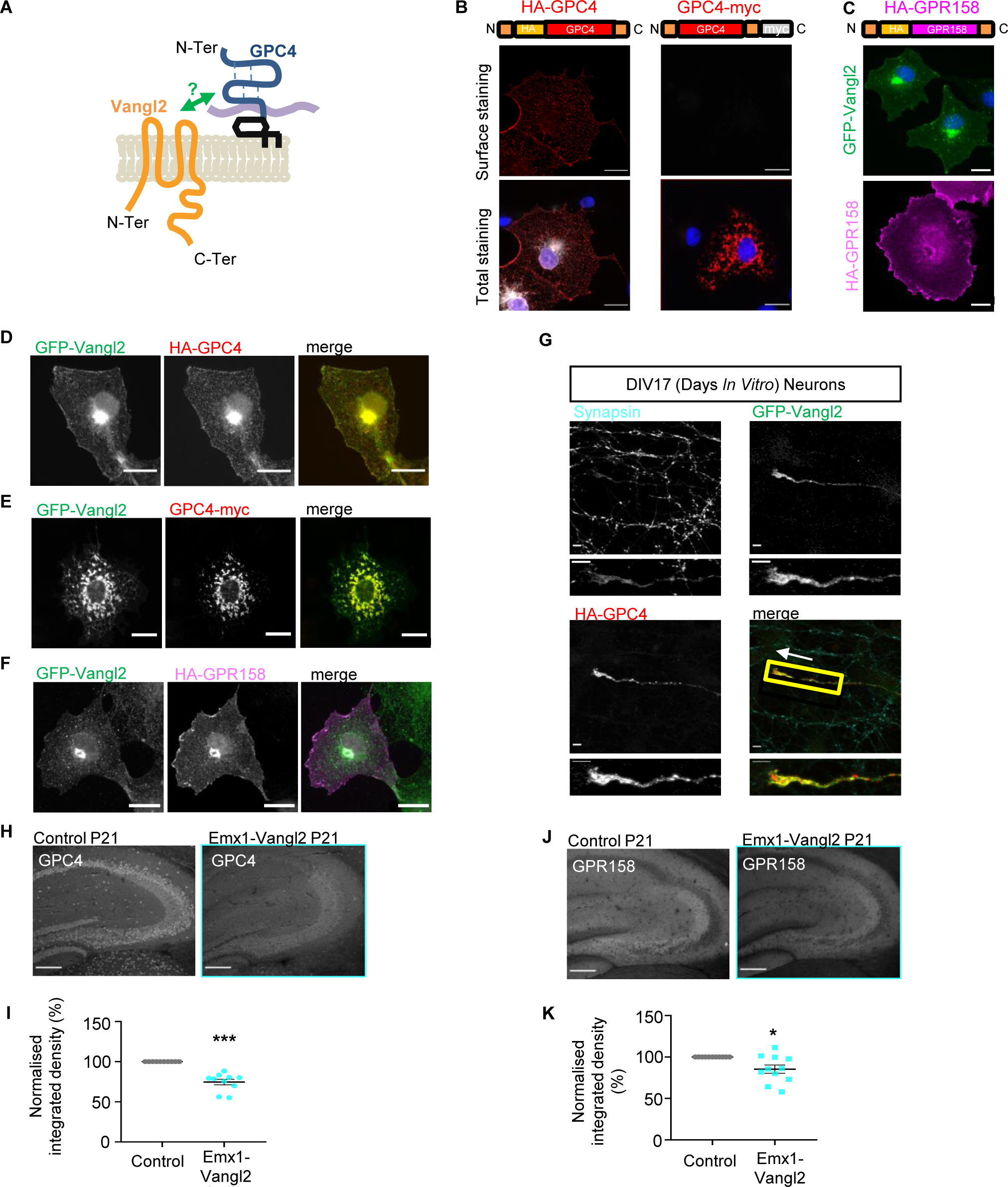
Vangl2 is associated with GPC4 in the presynaptic compartment early in development. **(A)** Schematic representation of Vangl2 and GPC4 protein structures and their potential interaction in *cis*. **(B)** Representative images of the surface and total labelling of HA-GPC4 (red) and GPC4-myc (grey) expression in transfected COS-7 cells. Scale bar, 20 µm. **(C)** Representative images of total GFP-Vangl2 (green) labelling and HA-GPR158 (magenta) expression in transfected COS-7 cells. Scale bar, 20 µm. **(D-F)** Representative images of total labelling of GFP-Vangl2 (green), HA-GPC4 (red), GPC4-myc (red) and HA-GPR158 (magenta) expression in co-transfected COS-7 cells. Scale bar, 20 µm. **(G)** Representative image of the expression of synapsin1 (blue), HA-GPC4 (red) and GFP-Vangl2 (green) in the axonal projection of a DIV17 transfected neuron. Scale bar, 5 µm. **(H-J)** Representative epifluorescence images of the *sl* with GPC4 (H) and GPR158 (J) labelling in P21 Emx1-Vangl2 and control mice. Scale bar, 200 µm. **(I-K)** Quantification of the percentage of reduction of the normalised integrated density for the signal of GPC4 (I) and GPR158 (K) in P21 control (grey) and Emx1-Vangl2 (turquoise) mice. N control = 11 mice, N Emx1-Vangl2 = 11 mice. One sample t-test ***P < 0.001 *P < 0.05. For I and K data are presented as mean ± SEM and single data points are shown as dot.

To further support these results, we examined the effect of early *vangl2* deletion on GPC4 levels at MfB/TE synapses. The integrated density of the fluorescent signal for GPC4 was significantly reduced in Emx1-Vangl2 mice compared to controls in 3-week-old mice hippocampal sections in the *sl* (−25.29±3.50%, one-sample t-test ***p<0.0001) (Figure 3H and 3I). As GPC4 binds in *trans* to the orphan receptor GPR158 to stabilise it at the postsynaptic side ^10^, we also quantified GPR158 immunofluorescence. As expected, GPR158 was significantly reduced in Emx1-Vangl2 mice compared to controls (−14.66±4.95%, One sample t-test *p=0.0143) (Figure 3J and 3K).

### Vangl2 is necessary for structural plasticity of adult MfB/TE synapses

To investigate whether the morphological deficits observed in young Emx1-Vangl2 mice were maintained in adult mice, we injected an AAV virus to re-express Vangl2 in 8-week-old mice and analysed MfB morphology at 10-week-old (Figure 4A). We observed no difference between genotypes in the morphological parameters of presynaptic MfBs in 10-week-old mice (Figure 4B). There was no statistical difference in volume (Figure 4C; control: 18.93±0.75 µm^3^, Emx1-Vangl2: 18.43±0.76 µm^3^; Mann-Whitney test ns p= 0.5702) or surface area (Figure 4D; control: 180.01±5.83 µm², Emx1-Vangl2: 181.62±5.71 µm²; Mann-Whitney test ns p= 0.6931). The number of filopodia was also unchanged between genotypes (Figure 4E; control: 0.63±0.06 filopodia, Emx1-Vangl2: 0.71±0.07 filopodia; Mann-Whitney test ns p= 0.5522), nor was the percentage of MfB with filopodia (Figure 4F; control: 40.38±3.37%, Emx1-Vangl2: 42.65±3.47%; Mann-Whitney test ns p= 0.6387). These results demonstrate that the early presynaptic morphological deficits observed at P25 are compensated in adult mice.

**Fig. 4:**
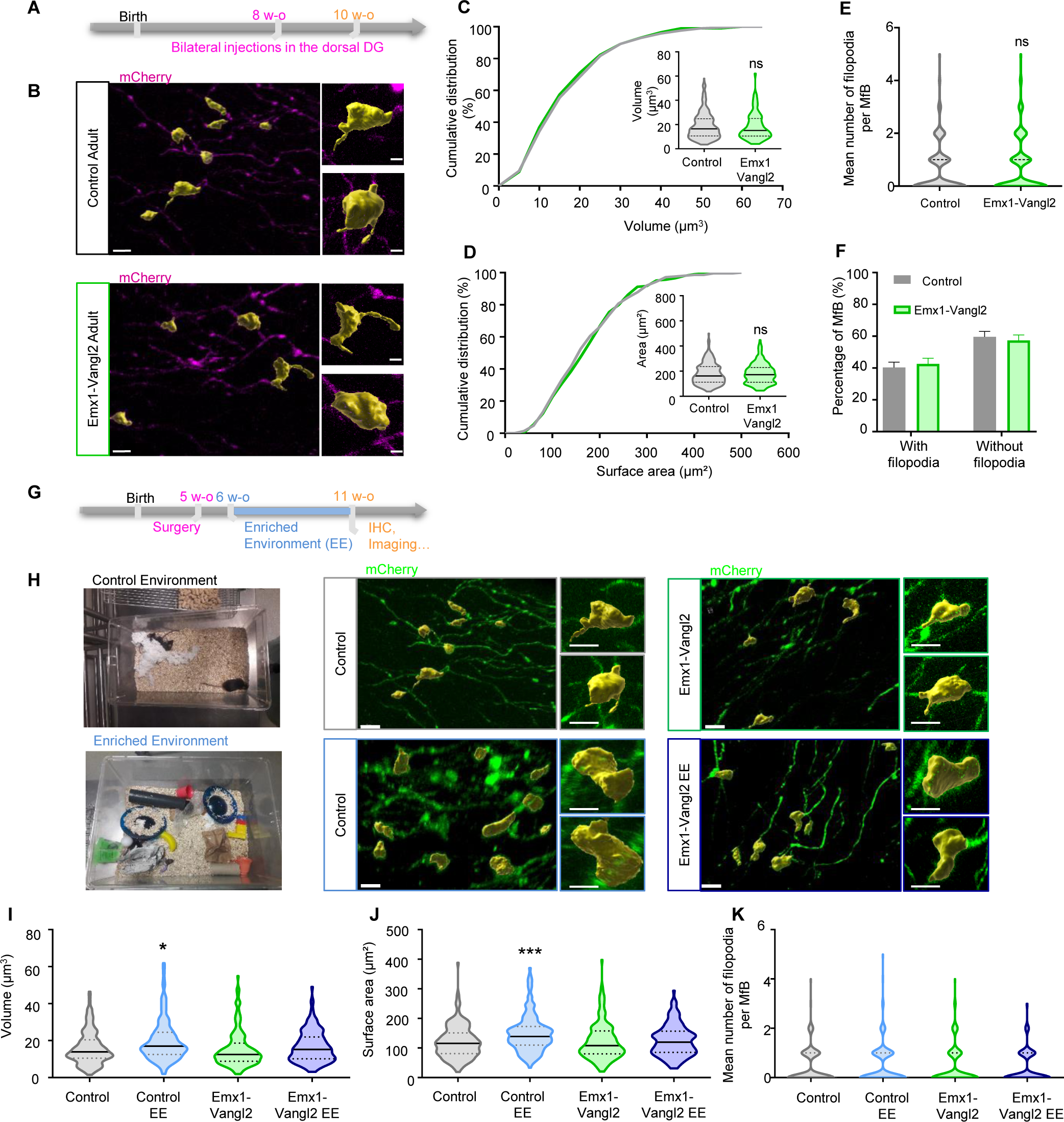
Morphological deficits of MfB are compensated in adult mice but their structural plasticity is still affected. (**A**) Experimental design for stereotaxic virus injections in the DG of 8-week-old mice. **(B)** Representative confocal images of labelled MfB (magenta) with 3D reconstruction (yellow) in the *sl* of 10-week-old Emx1-Vangl2 and control mice. Scale bar, 5µm. **(C-F)** Quantifications of volume (C), surface area (D), number of filopodia per MfB (E), and percentage of MfB with filopodia (F) in 10-week-old Emx1-Vangl2 and control mice. N control = 213 MfB from 6 mice, N Emx1-Vangl2 = 204 MfB from 6 mice. Mann-Whitney test, ns, p>0.05. **(G)** Schematic representation of the EE procedure in adult mice. To summarise, 5-week-old mice are injected with a mCherry expressing virus to label the MfB and then placed into an enriched (EE) or control (CE) environment for 5 weeks before doing immunohistochemistry and 3D confocal imaging of the MfB. **(E)** 3D reconstruction of presynaptic MfB (yellow) on confocal stacks in the four conditions: control (red) and Emx1-Vangl2 (blue) mice in EE, Control (grey) and Emx1-Vangl2 (green) mice in the CE. Scale bar, 5 µm. **(I-K)** Quantification of the MfB volume (I), surface area (J) and number of filopodia per MfB (K) for Emx1-Vangl2 and control mice after 5 weeks in EE or CE. N Control = 167 MfB from 7 mice; Emx1-Vangl2 = 167 MfB from 6 mice; Control EE = 177 MfB from 6 mice; Emx1-Vangl2 EE = 156 MfB from 6 mice. Kruskal Wallis test, *p<0.05, ***p<0,001. For C to D and I to K data are presented as median with 25th and 75th percentile. The shaded area represents the probability distribution of the variable. For F data are presented as mean ± SEM.

MfB/TE giant synapses are prone to morpho-functional plasticity, and mice exposed to an enriched environment (EE) respond by increasing the number and complexity of boutons and by stabilising their structures ^19,20^. To assess whether Vangl2 is necessary for this structural plasticity, we subjected Emx1-Vangl2 mice and littermate controls to an enriched or controlled environment for 5 weeks (Figure 4G and 4H). We validated the hypothesis as the volume and surface area of the MfB of control mice placed in EE compared to the controlled environment increased (Figure 4I and 4J; volume control: 16.01±0.6917 µm^3^, volume control EE: 19.80±0.8251 µm^3^, Kruskal-Wallis test *p<0.001; area control: 121.1±4.258 µm², area control EE: 149.4±4.697 µm², Kruskal-Wallis test ***p<0.0001). Notably, EE did not induce any change in the volume and surface area of the MfB in the absence of Vangl2 (Figure 4I and 4J; volume Emx1-Vangl2: 15.35±0.75 µm^3^, volume Emx1-Vangl2 EE: 16.71±0.67 µm^3^, Kruskal-Wallis test ns p>0.9999; area Emx1-Vangl2: 122.44±4.81 µm², area Emx1-Vangl2 EE: 125.06±4.17 µm², Kruskal-Wallis test ns p>0.9999). We observed no significant changes in the number of filopodia per synapse after exposure to EE, both in control and mutant mice (Figure 4K; control: 0.44±0.05 filopodia, control EE: 0.51±0.06 filopodia, Emx1-Vangl2: 0.43±0.05 filopodia, Emx1-Vangl2 EE: 0.39±0.05 filopodia; Kruskal-Wallis test ns p=0.53). These results demonstrate that, although the effect of early loss of *Vangl2* is morphologically compensated in adult mice, the structural plasticity of MfB/TE synapses is impaired in the mutants.

### Defects in MfB/TE synapse morphology are accompanied by defects in synaptic transmission and plasticity

To investigate the effect of Vangl2 loss on synaptic transmission at MfB/TE synapses, we first recorded CA3 PCs in acute hippocampal slices from P21-P32 Emx1-Vangl2 mice and control littermates in whole-cell voltage-clamp configuration. The frequency of spontaneous excitatory postsynaptic currents (sEPSC) was reduced by 50% in Emx1-Vangl2 mice compared to control (Figure 5A to 5C; control: 4.0±0.47 Hz, Emx1-Vangl2: 2.02±0.25 Hz; unpaired t-test ***p=0.0007). The amplitude of sEPSCs was also reduced in Emx1-Vangl2 mice (Figure 5D and 5E; control: 25.57±1.34 pA, Emx1-Vangl2: 21.03±0.84 pA; unpaired t-test **p=0.0068). A detailed comparative analysis of the distribution of sEPSC amplitudes showed a significant decrease in synaptic events with amplitudes above 30 pA in Emx1-Vangl2 mice (Figure S4A to S4C). These results show that Vangl2 loss results in a specific impairment of larger sEPSC amplitudes at P21 MfB/TE synapses ^10,21,22^. Next, we investigated whether Vangl2 loss affects the glutamate receptor composition of MfB/TE synapses. We recorded AMPA/Kainate receptor (hereafter AMPAR) EPSCs and NMDA receptor (NMDAR) EPSCs evoked by Mf stimulation in acute hippocampal slices from Emx1-Vangl2 mice and control littermates. There was no significant difference in AMPAR and NMDAR EPSC amplitudes (Figure 5F to 5K). Thus, the functionality of postsynaptic receptors at MfB/TE synapses is not affected by the absence of Vangl2. Together, these results on basal transmission suggest a presynaptic function impairment in the absence of Vangl2, that could be consistent with a possible reduction of presynaptic release sites (active zones) per bouton, in the smaller MfBs observed in Emx1-Vangl2 mice (Figure 1E and 4B).

**Fig. 5:**
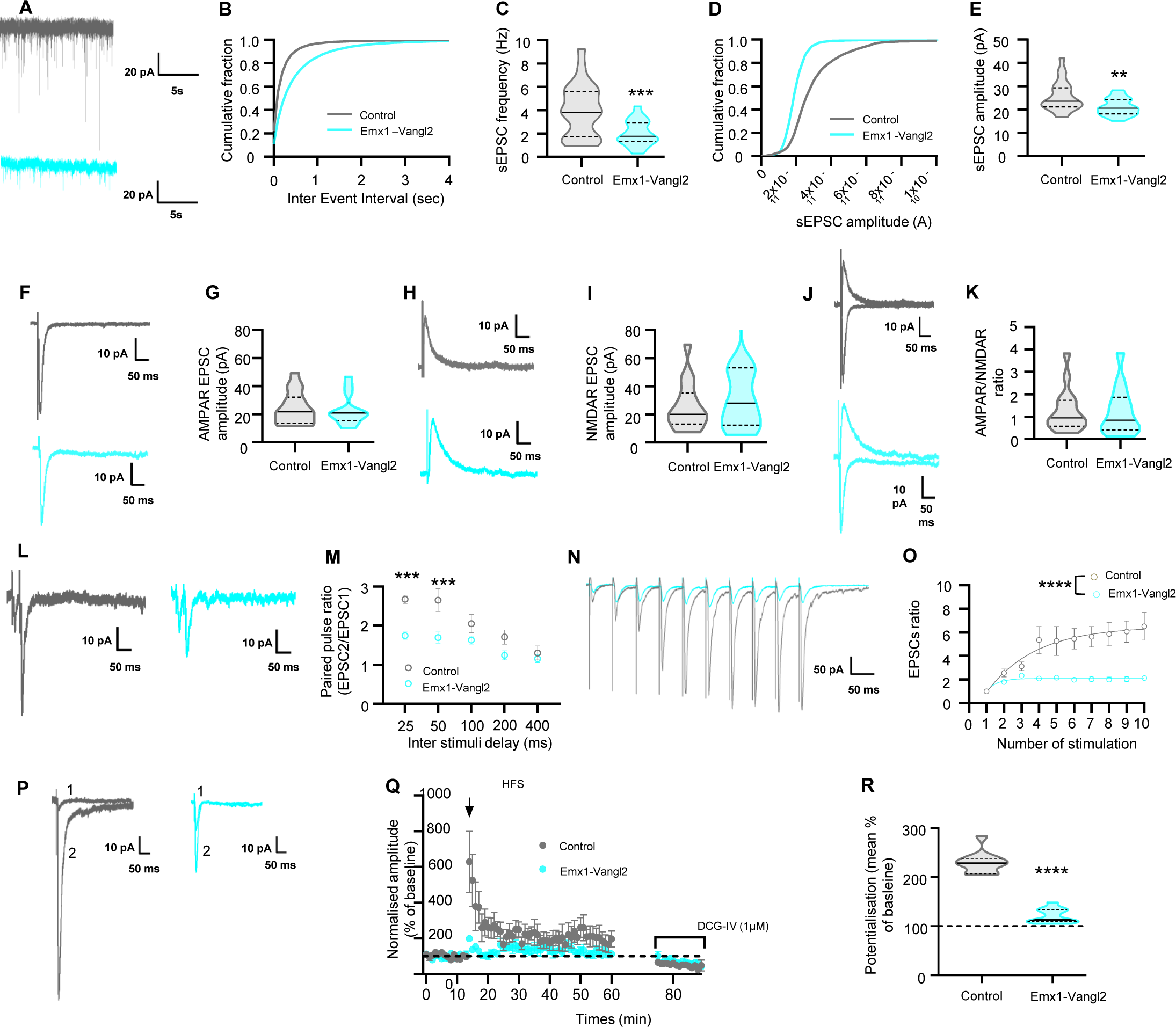
Emx1-Vangl2 mice exhibited reduced synaptic transmission and synaptic plasticity at MfB/TE synapses. **(A)** Representative spontaneous EPSC (sEPSC) traces from CA3a PC in P21-P32 control (grey) and Emx1-Vangl2 (turquoise) acute hippocampal slices. **(B, D)** Cumulative distribution of sEPSC inter-event intervals (B) and amplitudes (D) in control and Emx1-Vangl2 neurons. **(C, E)** Quantification of sEPSC frequency and amplitude of control neurons (N = 16 mice, n = 24 cells) versus Emx1-Vangl2 (N = 14 mice, n = 20 cells). ****p<0.0001, **p = 0.0036. Unpaired t-tests with Welch’s correction. **(F, H)** Representative traces of AMPA/Kainate receptor evoked responses (F) and evoked response of NMDA receptors from CA3a PC (H) in P21-P32 control and Emx1-Vangl2. **(G, I)** Quantification of AMPA (G) and NMDA (I) receptors EPSC amplitudes of MfB/TE synapses in control (N =12 mice, n =16 cells) and Emx1-Vangl2 mice (N = 11 mice, n =13 cells). ns, p>0.05 **(J) Representative** traces of AMPAR/NMDAR ratio from CA3a PC in P21-P32 control and Emx1-Vangl2. **(K)** AMPAR/NMDAR ratio at MfB/TE synapses in control (N = 12 mice, n = 16 cells) Emx1-Vangl2 mice (N = 11 mice, n = 13 cells). ns, p>0.05. **(L, M)** Representative traces of PPR and quantification of PPR of MfB/TE synapses in control (N = 13 mice, n = 14 cells) and Emx1-Vangl2 mice (N = 12 mice, n = 15 cells). Genotype effect and ISI effect ***p<0.001, two-way ANOVA with Bonferroni’s multiple comparisons test. **(N)** Representative traces of train of 10 stimuli at 20 Hz of MfB/TE synapses of control and Emx1-Vangl2 mice. **(O)** Analysis of the presynaptic facilitation in trains of 10 stimuli at 20Hz in control and Emx1-Vangl2 MfB/TE synapses. Two-way ANOVA; control: N = 14 mice, n = 16 cells, Emx1-Vangl2: N = 9 mice, n =12 cells; genotype effect **** p<0.0001. **(P)** Representative traces of EPSC before (1) and after (2) HFS protocol of MfB/TE synapses of control and Emx1-Vangl2 mice. **(Q)** Summary graphs of LTP time course recorded over 50 minutes in control (N=6 mice, n=6 cells) and Emx1-Vangl2 mice (N = 4 mice, n = 7 cells). **(R)** Analysis of LTP magnitude 40-50 minutes after the induction of tetanus in control and Emx1-Vangl2. (Unpaired t-tests with Welch’s correction; control: N = 6 mice; n = 6 cells, Emx1-Vangl2: N = 4 mice, n = 7 cells, ****p<0.0001). For C, E, G, I K and R data are presented as median with 25th and 75th percentile. The shaded area represents the probability distribution of the variable. For M, O and Q data are presented as mean ± SEM.

To determine the impact of the loss of Vangl2 in presynaptic release at MfB/TE synapses, we analysed paired-pulse facilitation (PPF) (Figure 5L and 5M). With inter-stimulus intervals (ISI) of 25 ms and 50 ms, the paired-pulse ratio (PPR) was markedly decreased in Emx1-Vangl2 mice (25 ms: PPR = 174% Emx1-Vangl2 mice as compared to PPR = 267% in control mice; 50 ms, PPR = 169% in Emx1-Vangl2 mice as compared to PPR = 265% in control mice, two-way ANOVA, ISI and genotype effects: ***p<0.001). No significant difference was observed at higher inter-stimulus intervals (Figure 5M). We also analysed presynaptic facilitation in trains of 10 stimuli at 20 Hz in control and Emx1-Vangl2 MfB/TE synapses (Figure 5N and 5O). Presynaptic facilitation was almost abolished in Emx1-Vangl2 MfB/TE synapses (two-way ANOVA, stimulation number and genotype effects ****p<0.0001).

We also investigated long-term potentiation (LTP) which depends on presynaptic mechanisms at MfB/TE synapses. A high-frequency stimulation (HFS) protocol elicited robust LTP of EPSCs in control synapses that lasted for at least 45 minutes (Figure 5P and 5Q). The magnitude of LTP was significantly reduced in Emx1-Vangl2 synapses (Figure 5R; control: 229.2±6.70%; Emx1-Vangl2: 120.9±4.41%; Mann-Whitney test *** *p<0.0001). The level of EPSCs potentiation measured immediately after HFS was also altered in Emx1-Vangl2 synapses as compared to control synapses (control: 629.1.10±172.7%; Emx1-Vangl2: 199.00±16.86%), suggesting that the two groups were not similarly activated by HFS (Figure 5Q). Collectively, these results indicate that *Vangl2* deletion impairs synaptic transmission, short-term and long-term synaptic plasticity predominantly on the presynaptic side of MfB/TE synapses in the CA3a region of the hippocampus.

### Vangl2 is necessary for the flexibility of relational/declarative memory

To determine the functional consequences of the early absence of Vangl2 in the forebrain, we tested 10-week-old Emx1-Vangl2 mice and their littermate controls in a series of tasks measuring emotional, exploratory, and cognitive behaviour. We first performed elevated plus-maze (Figure S5A) and dark/light experiments (Figure S5C) to test anxiety-like behaviour. Emx1-Vangl2 mice showed no change in anxiety-like behaviour (Figure S5B and S5D) and locomotion activity and exploration were also unaltered (Figure S5E to S5G). We next analysed the effect of Vangl2 loss on spatial learning and memory in the Morris water maze (MWM), a task which is known to depend on the hippocampus. We found no impairment in the Emx1-Vangl2 mice, which were indistinguishable from their controls in learning with visual or spatial cues and in reference memory in the MWM task (Figure S5H to S5K).

Next, we tested another aspect of spatial reference memory: its flexibility, known to rely on relational representation and taken as a model of declarative memory, using a specific version of the 8-arm radial maze task (Figure 6A) ^23–25^. The acquisition phase of this task assesses the ability to learn constant food locations within 3 invariant pairs of adjacent arms. In this phase, we found no impairment in Emx1-Vangl2 mice, as they reached the learning criterion as quickly as control mice (t-test, t24=1.4) (Figure 6B) and had similar levels of performance during the first five days of training (Figure 6C; Table S1) and the last five days of training (Figure 6D; Table S1). Both groups were equally good at acquiring spatial reference memory. The test phase then assesses the ability to flexibly express learned information in a modified situation, a characteristic of declarative memory. This phase revealed an impairment of memory flexibility in Emx1-Vangl2 mice. We compared the performance on the last day of acquisition (D-1) with the performance on the test day. There was no difference between D-1 and the test regardless of genotype with respect to the unmodified control pair (Figure 6E: pair C; Table S1) or the novel pair (i.e., unlearned control pair N) for which test performance was compared to chance level (Figure 6F; Table S1). In the critical flexibility test (recombined pair AB made of arms that were featured in separate pairs in the acquisition), the performance declined significantly between the end of acquisition D-1 and the test only in Emx1-Vangl2 mice (Figure 6G; Table S1). Thus, while control mice were able to flexibly express their memories in a modified test situation, the Emx1-Vangl2 mice failed to do so. These data demonstrate that Emx1-Vangl2 mice can learn and express spatial memories in routine situations, but they are unable to adapt their spatial memory in a changing situation, indicating memory inflexibility.

**Fig. 6:**
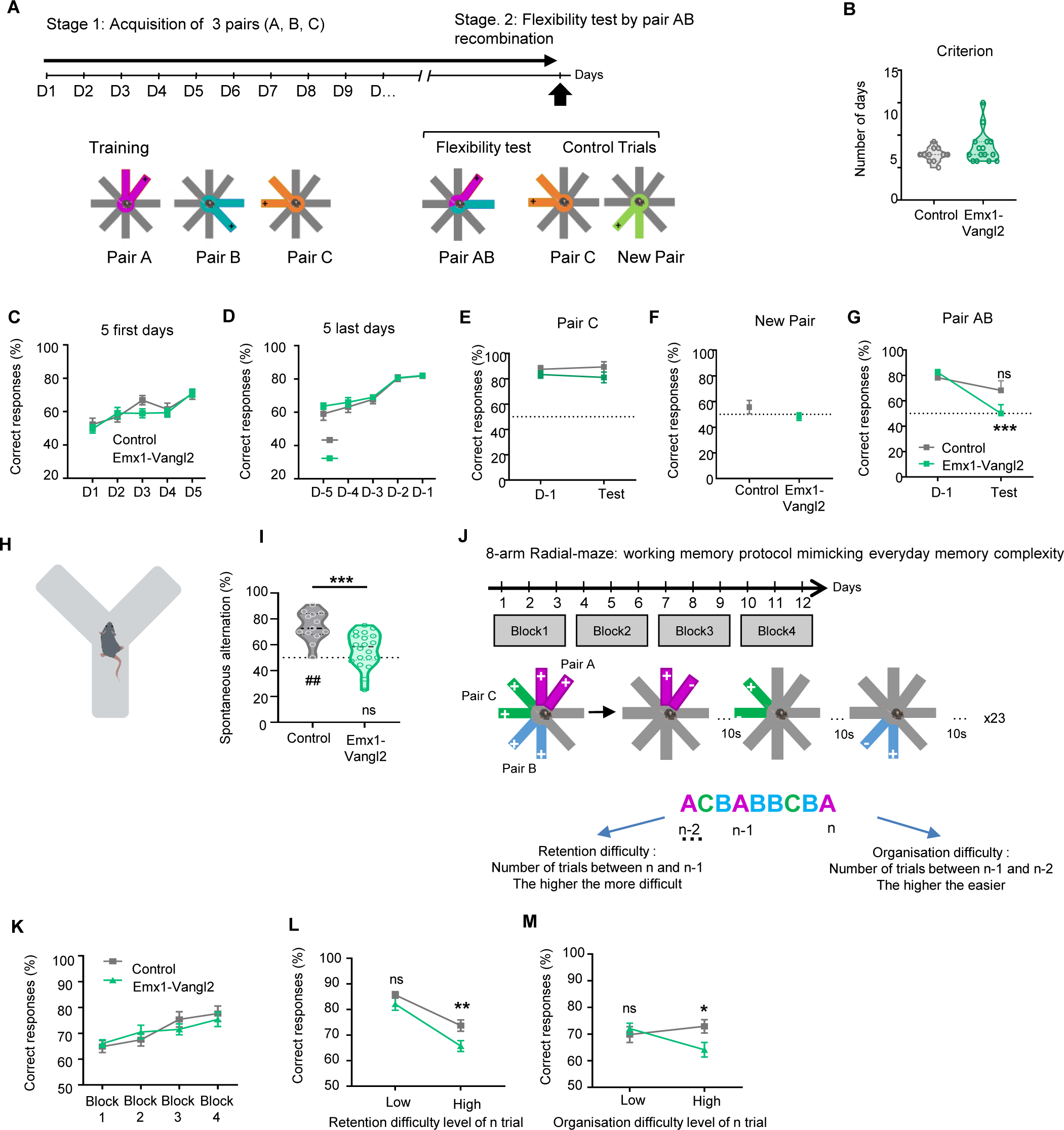
Early *vangl2* deletion impairs the flexibility of spatial reference memory and working memory in complex memory tasks in adult mice. **(A)** Experimental design for the 8-arm radial maze (RM) test reference memory. **(B-D)** Quantification of the performance level during the acquisition phase is measured by the number of days before reaching criterion (B) and the percentage of correct responses during the 5 first (C) and 5 last days of acquisition before the reaching of the learning criterion(D). N: Control = 11, Emx1-Vangl2 = 15. Mann-Whitney test and two-way ANOVA. **(E-G)** Evolution of the percentage of correct responses between the last day of acquisition (D-1) and the test for the control pairs (E-F) (Pair C is a “known” control; New Pair is a “novel” control appearing in Test only) and critical pairs (Pairs A and B recombined into Pair AB in the Test, representing the critical flexibility probe) (G). N Control = 11, N Emx1-Vangl2 = 15. For the Novel Pair: Mann-Whitney and One Sample t-test vs 50%. For Pair C and Pair AB: two-way ANOVA with Bonferroni’s comparison, D-1 versus Test, ns, p>0.05; *** p<0.001. **(H)** Experimental setup for the spontaneous alternation test in Y maze. **(I)** Mean percentage of spontaneous alternation control (grey) and Emx1-Vangl2 adult mice (green). N Control = 13 mice; N Emx1-Vangl2 = 20 mice. Unpaired t-test, ***p<0.05; One sample t-test against 50%, ##p<0.001, ns, p>0.05. **(J)** Experimental design for the working memory in the 8-arm radial maze test. **(K)** Quantifications of the percentage of correct response during the first four blocks of the experiment (ITI 10s). N Control = 12, N Emx1-Vangl2 = 13. Two-way ANOVA, ns, p>0.05. **(L-M)** Quantifications of the percentage of correct responses depending on the level of retention (L) and organisation (M) difficulty. N Control = 12, N Emx1-Vangl2 = 13. Two-way ANOVA, *p<0.05, ***p<0.001. For B and I data are presented as median with 25th and 75th percentile and single data points are shown as dot. The shaded area represents the probability distribution of the variable. From C to G and K to M data are presented as mean ± SEM.

### Vangl2 is necessary for both retention and organisation of working/everyday-like memory

Next, we investigated spatial working memory, which, unlike reference memory, is constantly changing. First, we used a simple test based on the natural tendency of mice to explore a novel environment, i.e., spontaneous alternation in the Y-maze (Figure 6H). The performance of Emx1-Vangl2 mice was not significantly different from chance, whereas the control group had an alternation score of almost 75% (Figure 6I; Table S1). Since Y-maze alternation requires short-term memory of the previously visited arm, these data suggest that Emx1-Vangl2 mice have poor spatial working memory. To confirm this suggestion, we assessed spatial working memory in another radial-maze task that mimics the complexity of memory in everyday life ^26^ (Figure 6J). The task consists of the repeated use of 3 pairs of adjacent arms (pairs A, B and C), with the location of the food varying within each pair according to an alternation rule: the baited arm in each (n) trial is always the one that was not visited by the animal in the previous trial (n-1) with the same pair. We can test two memory components simultaneously, retention and organisation, since it is necessary to continuously retain (not just 1, but 3) previously visited arms (one for each pair) and to organise/update these memories so as not to mix up the pairs and successive trials. The animals were subjected to 12 days of training, and both groups had a similar progression in the percentage of correct responses over the four 3-day blocks of training (Figure 6K; Table S1). The differences between genotypes emerged when performance in the last two blocks were analysed as a function of either ‘retention difficulty’ (Figure 6L) or “organisation difficulty” (Figure 6M). As expected, when the trials were divided into two levels of retention difficulty (low vs. high) we observed that performance decreased with increasing retention difficulty in both groups. However, Emx1-Vangl2 mice had significantly lower performance than controls only at high retention difficulty (Figure 6L; Table S1). When the trials were divided into two levels of organisation difficulty, Emx1-Vangl2 mice performance dropped significantly as the difficulty of the organisation increased, whereas the performance of controls remained stable across the two difficulty levels. (Figure 6M; Table S1). These results show that Emx1-Vangl2 mice have specific impairments in both the retention and organisation components of everyday-like / working memory.

### Vangl2 contributes to contextual fear memory formation and its absence induces maladaptive memory for salient but irrelevant cues

To assess the type of cues the animals use to recall information, we used a contextual fear conditioning paradigm using a tone-shock unpairing procedure where only the context (the chamber) is predictive of the footshock, not the tone (see Methods). During the conditioning phase, the mice are placed on a footshock grid in a translucent cage providing visual access to the surrounding experimental room, and presented with a non-predictive tone that is pseudorandomly distributed relative to the shocks. This tone-shock unpairing procedure allowed us to assess the hippocampus-dependent processing of distal and tonic contextual cues (vs. proximal and/or phasic, salient cues) allowing the formation of a complex representation of the environment (Figure 7A). During the first auditory cue test, the mice are placed in a neutral (and opaque) chamber and presented with the irrelevant (not predictive) tone. The level of freezing was low in both control and Emx1-Vangl2 mice during the first two minutes, as the neutral chamber is not associated with the shock. At the onset of the tone, freezing levels were still low in both genotypes. However, compared to control mice, Emx1-Vangl2 mice displayed an increase of their freezing responses after the end of the tone, which indicates an abnormal cue-based association between the non-predictive salient tone and the shock (Figure 7B; Table S1).

**Fig. 7:**
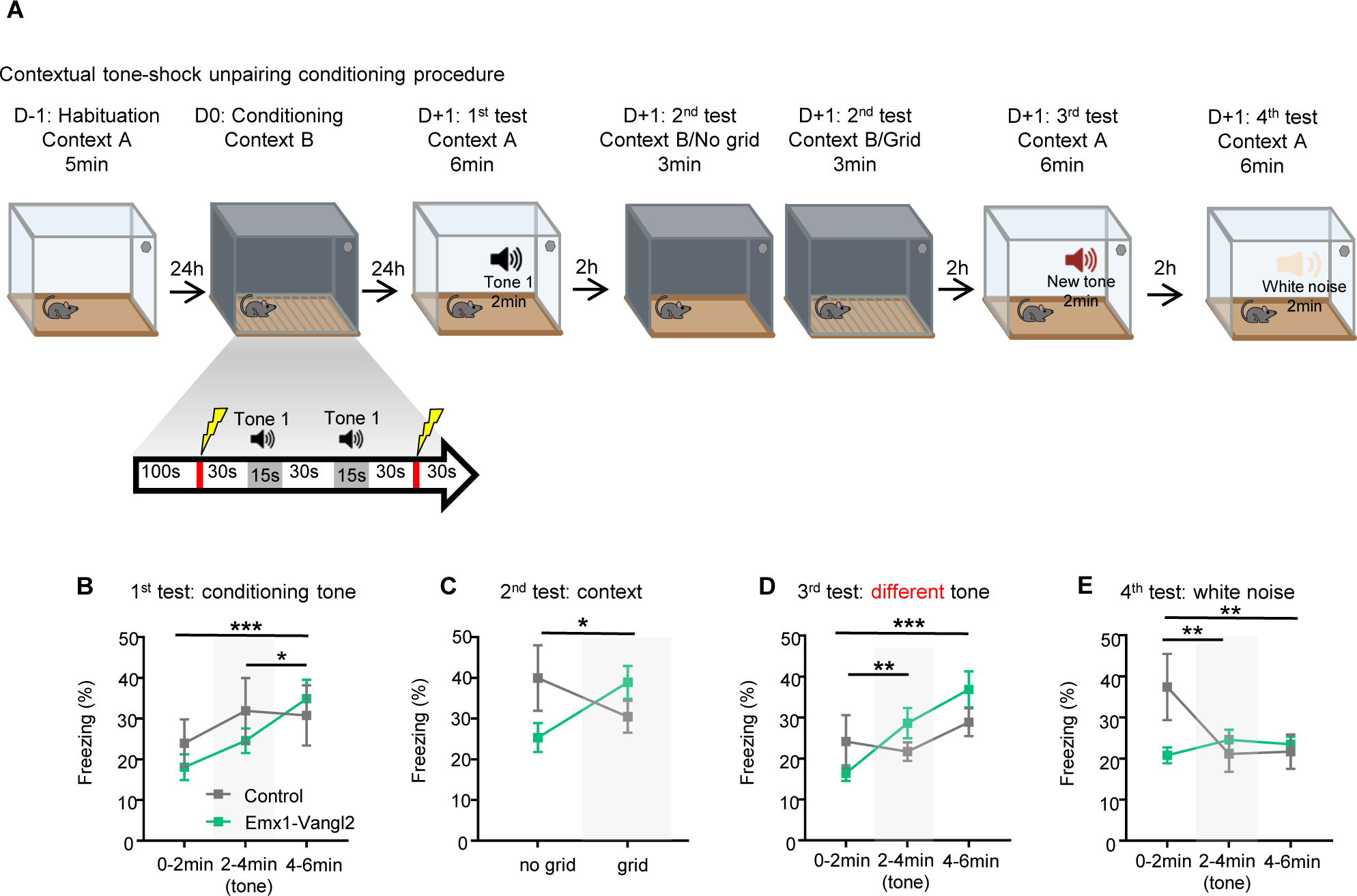
Emx1-Vangl2 mice show an imbalance between contextual conditioning and cued conditioning. **(A)** Experimental design for the contextual fear conditioning protocol with tone-shock unpairing. **(B)** Quantification of the percentage of freezing during the first conditioning tone test. N Control = 7, N Emx1-Vangl2 = 20. Two-way ANOVA, *p<0.05 *** p<0.001. **(C)** Quantification of the percentage of freezing during the contextual test. N Control = 7, N Emx1-Vangl2 = 20. Two-way ANOVA, *p<0.05. **(D)** Quantification of the percentage of freezing during the partial generalisation test. N Control = 7, N Emx1-Vangl2 = 20. Two-way ANOVA, **p<0.01 *** p<0.001. **(E)** Quantification of the percentage of freezing during the global generalisation test. N Control = 7, N Emx1-Vangl2 = 20. Two-way ANOVA, **p<0.01. For B to E data are presented as mean ± SEM.

The second test assessed contextual conditioning, comparing distal (visuo-spatial cues in the room) vs proximal (the grid) contextual cues. When the electric grid was hidden, control mice displayed higher freezing than Emx1-Vangl2 mice indicating an adaptive contextual fear memory based on the hippocampus-dependent processing of distal contextual cues (Figure 7C; Table S1). In contrast, when the platform is removed, then revealing the grid (3min later), while controls displayed a decrease in their freezing responses (due to intra-trial extinction), Emx1-Vangl2 mice displayed an increase in their fear responses indicative of a fear memory mainly based on proximal salient cues, a simple form of associative memory which does not require the integrity of the hippocampus (Figure 7C; Table S1). The third test used a similar but different tone from the one used in the conditioning phase to assess a possible partial generalisation of fear (Figure 7A). Compared to control mice, Emx1-Vangl2 mice displayed an increase in their freezing levels, indicating a partial generalisation of the fear response to a similar salient simple cue (Figure 7D; Table S1). Finally, during the fourth test in the neutral chamber, the mice were exposed to a white noise to assess a possible global generalisation of fear (Figure 7A). Before the white noise was presented, the freezing level of the Emx1-Vangl2 mice was low, while that of one of the control mice was unexpectedly high (Figure 7E; Table S1). This relatively high fear response in controls may indicate a loss of context specificity but also reflects to some extent a certain expectation regarding the possible occurrence of the shock at any time in this type of conditioning procedure in which the shock is not temporally predicted by the occurrence of a phasic cue (tone-shock unpairing procedure). When the white noise started, the freezing level of the control mice decreased. Freezing in Emx1-Vangl2 mice remained low in both genotypes after the tone ended, indicating that there was no association between the white noise and the footshock, and therefore no global generalisation of fear (Figure 7E; Table S1). Altogether, these results indicate that Emx1-Vangl2 mice displayed an abnormal fear memory mainly based on the processing of salient proximal and tonic (grid) and/or salient phasic (tone) cues, whatever their relevance in the learning procedure, rather than (predictive) distal contextual cues.

## Discussion

Our data show that Vangl2 is required for the morpho-functional development and maintenance of MfB/TE synapses. Its absence has long-lasting consequences on synapse physiology and the hippocampal circuitry, with selective plasticity and memory deficits in adult mice. This establishes Vangl2 as a key regulator of DG-CA3 connectivity and functions during postnatal development and adulthood.

Using 3D reconstructions of MfB/TE synapses, we show that early *vangl2* deletion delays synapse morphogenesis, with smaller and less complex immature-like synapses at P21. The spatio-temporal profile of Vangl2 expression in the *stratum lucidum* of CA3 from P7 to P21 is consistent with a function as a critical modulator of MfB/TE synapse morphogenesis during development. The expression of Vangl2 in the GCs of the DG ^13^, together with its presence in the growth cone of cultured hippocampal neurons (this study), supports a presynaptic function of the protein at MfB/TE synapses ^15^. Other studies have reported a postsynaptic localisation of Vangl2 in brain samples ^27^ or in the spines of CA1 pyramidal neurons and DG granule cells *in vivo* ^13,28^. The subcellular localisation of Vangl2 is likely to be region, if not synapse, specific. We propose that at MfB/TE synapses Vangl2 is present in both the pre- and postsynaptic compartments.

The role of Vangl2 in the regulation of tissue and cell morphogenesis, including in subregions of the hippocampus, is not unexpected. PCP or Wnt/PCP signaling is a widely conserved pathway originally known to coordinate cell orientation within the plane of an epithelium ^29^. PCP is regulated by evolutionarily conserved proteins, of which Vangl2 is the bone fide protein. In vertebrates and mammals, the Wnt/PCP signaling pathway controls various morphogenetic processes in most tissues and provides positional cues to cells for directed movements, particularly during gastrulation and neurulation ^30^. Mutations in Vangl2 are associated with severe disorders of the nervous system called neural tube defects (NTDs), such as spina bifida and anencephaly ^31^. During these embryonic processes, Vangl2 is thought to regulate Wnt/PCP signaling in a dosage-sensitive manner, including at the local level ^32^. We found that the levels of the HSPG protein GPC4 were reduced in the Emx1-Vangl2 mutant. GPC4 is a heparan sulphate proteoglycan that promotes Wnt/PCP signaling ^18^. Genetic epistasis experiments in zebrafish have identified functional interactions between GPC4 and Vangl2 ^33^, while in zebrafish endodermal cells, GPC4 deficiency is thought to disrupt polarised Wnt-PCP signaling by interfering with the asymmetric accumulation of Vangl2 ^18^. GPC4 could act as a co-receptor or modulator of Vangl2 in the presynaptic bouton of MfB/TE synapses, participating in its stabilisation at the membrane via its GAG domains ^34^ or simply by modulating its interaction with other proteins ^35^. The absence of Vangl2 in the mutant could destabilise the known transsynaptic interaction between GPC4 and GPR158, disrupting the development of MfB/TE synapses.

The alteration of the putative Vangl2-GPC4 interaction could also have morphological consequences via their common interactor N-cadherin ^18^. Our group and others have shown that Vangl2 is involved in N-cadherin turnover, in the growth cone of hippocampal neurons regulating axonal outgrowth, and in spine formation ^14,15^. N-cadherin plays a critical role in the stabilisation and increased complexity of the MfB and TE ^36^. Also, alteration of N-cadherin dynamics could induce a more general reduction of synaptic vesicle turnover and of the number of vesicles in the reserve pool ^37,38^. We observed reduced molecular levels of synaptic proteins synaptoporin and synapsin1 in Emx1-Vangl2 mutant mice, which are consistent with altered trafficking and docking mechanisms required for neurotransmitter release ^39,40^. The absence of Vangl2 is therefore likely to destabilize N-cadherin-dependent mechanisms, resulting in both morphological and functional deficits of MfB/TE synapses.

We found that spontaneous synaptic transmission activity and synaptic plasticity were altered in 3-week-old mice without Vangl2. However, we did not find any alteration in the AMPAR/NMDAR ratio in the absence of Vangl2. It is likely that the reduced surface and volume of MfB in the absence of Vangl2 goes along with a decrease in the number of releasing sites for synaptic vesicles (active zones) per bouton, thus decreasing the frequency of synaptic events, especially those of high amplitude, characteristic of this synapse. This is consistent with previous studies showing that MfB size correlates with the number of active zones ^41,42^ supporting a presynaptic MfB deficit in early postnatal stages. It is also possible that changes in the postsynaptic response occur later in development. This might lead to functional maturation defects at an older age. Such phenotypes have been observed in GluK2 mutants, which also show defects in synaptic morphogenesis ^1^.

It has been shown that spontaneous and evoked release occurring at the same synapse in the hippocampus involve distincts pools of AMPA and NMDA receptors ^41^. This could also explain why we observe only spontaneous synaptic transmission impairment in Emx1-Vangl2 mice. This group also suggested the existence of two different pools of vesicles involved in spontaneous and evoked release, with independent recycling mechanisms ^42^. Thus, the absence of Vangl2 migth only affect the recycling of spontaneous pool of vesicles. Moreover, there might be a distinct fusion machinery for spontaneous and evoked release ^43^. Although spontaneous and evoked release depend on the same SNARE machinery, the molecular interactions of these various SNARE components may be distinct. The absence of Vangl2 might impair synaptic vesicle fusion complex interaction and/or docking complex interaction as it has been mentioned previously with the alteration of the GPC4 and GPR158 interaction.

The morphological deficits observed earlier in the presynaptic bouton are no longer observed in adults. Thus, the early postnatal profile of expression of Vangl2 in the *sl* of CA3 is critical for morphological processes, but the absence of Vangl2 is later compensated. This compensatory mechanism could, at least in part, be due to other PCP members, including Vangl1. A similar delayed morphological compensation has been observed in the cochlear system that is highly dependent on Vangl2 for its morphogenesis ^29,44^. Dean and coworkers showed that loss of Vangl2 disrupts the planar orientation of the auditory stereociliary bundles in neonatal mice (P0-P3), but this misorientation is corrected in P10-P12 mice ^44^. Similar to the current study, the morphological compensation is not accompanied by functional compensation, as the hearing of *vangl2* mutants is still defective.

Impaired structural plasticity at MfB/TE synapses of adult Emx1-Vangl2 mice exposed to EE clearly demonstrated that Vangl2 is involved in adult structural plasticity. The change in structural plasticity is likely to also involve a change in synaptic transmission. Blocking neurotransmitter release via mGlu2 receptor activation blocks structural plasticity of MfB/TE synapses ^19,45^. LTP also induces MfB/TE synapse structural remodeling ^46^. The LTP deficit we observed in young mutant mice may have long-lasting consequences on the plasticity of MfB/TE synapses, both structurally and functionally, leading to permanent dysfunction of DG-CA3 circuits in Emx1-Vangl2 mice.

Early deletion of *vangl2* has long-lasting effects on some hippocampal-dependent processes, such as the flexibility of spatial reference memory, proposed as a model of declarative memory and both retention and organisational processes of complex working memory proposed as a model of everyday-like memory, while leaving spatial learning and recall in the MWM paradigm intact. The absence of alterations in spatial memory retrieval during MWM is consistent with studies in CA3-lesioned rats showing that alterations of MfB/TE synapses had no appreciable effect on this process ^47^. However, in an RM paradigm, Emx1-Vangl2 mutant mice showed a deficit in spatial reference memory flexibility, supporting a role for DG-CA3 connectivity in flexibility processes. This is consistent with studies reporting a strong involvement of DG GCs in tasks involving spatial flexibility, or studies linking increased adult neurogenesis to improved DG-CA3 activity and cognitive flexibility ^48–50^. The early change in the DG-CA3 network in the absence of Vangl2 would alter the encoding of new contextual memories and the adaptation of learned spatial information, leading to a “rigidity” of spatial reference memory.

Our Y-maze results, confirmed with the everyday like working memory task in the radial maze, show that our mice have a deficit of working memory in the absence of Vangl2 ^51^. Concerning the radial maze task, we observed a performance deficit in the mutant mice specifically when the retention and organisation demands are high. This indicates that early deletion of *vangl2* induces specific deficits in spatial working memory. This task requires the activation of the DG-CA3 circuit ^26^, while other working memory tests in the radial maze have shown the involvement of the dorsal DG ^52^. In particular, the test depends on 1) the effective encoding of different pairs based on DG-dependent pattern separation; 2) a combination of both pattern completion and separation for retrieval of the pairs. Moreover, the subregions CA3a and CA3b are involved in the encoding and retrieval of spatial working memory ^53^. The results of the fear conditioning experiments showed that the Emx1-Vangl2 mice displayed an impaired contextual conditioning, with the shock being abnormally associated with salient (but potentially irrelevant, e.g. non-predictive tone) cues. The deficits we observed in the *sl* of CA3 may be sufficient to explain the reduced retrieval of contextual cues and the abnormal generalisation of fear to non-predictive cues. Indeed, CA3 has been associated with the encoding and retention of contextual cues, whilst CA1 and the amygdala are involved in encoding and retrieval of both tonic and contextual cues ^54^. The DG-CA3 alterations in our mutant might be sufficient to induce an imbalance between contextual and cued conditioning.

We show that the early deletion of *vangl2* in the Emx1-Vangl2 mice has long-term consequences on the function of MfB/TE synapses, leading to disruption of DG-CA3 circuit function and a variety of impairments in hippocampal-dependent tasks. We do not exclude that other parts of the hippocampal circuitry - or beyond the hippocampal circuitry - are affected and contribute to the cognitive deficits we report ^31,55^. Nevertheless, this work and our previous study ^13^ demonstrate the importance of Vangl2 in the architecture and function of DG-CA3 connectivity involved in declarative memory tasks.

## Materials and Methods

### Animals

All procedures involving animals were performed in accordance with the European Union Directives (2010/63/EU) and the University of Bordeaux Ethical Committee (CE50) and French research Ministry according to the following three numbers: Dir13115, APAFIS #19761 and #27115. All the animals used in this study were generated in our breeding facility (accreditation number B33-063-090). The conditional knockout line (Emx1-Vangl2) was generated by crossing Vangl2 flox/flox animals ^56^ with Emx1B6.129S2-Emx1tm1(cre)Krj/J (Emx1-Cre) mice ^57^. Mice expressing the Cre recombinase and therefore lacking the expression of *Vangl2* were considered as our Emx1-Vangl2 mutant mice while their littermates which did not expressed the Cre recombinase were used as controls. Mice were maintained under standard conditions (food and water ad libitum; 23±1°C, 7-12 h light/dark cycle, light on at 7:00; experiments performed between 9:00 and 17:00). Male mice between 8 and 12 weeks of age were used, unless otherwise specified. Five to eight mice were housed together in polypropylene cages for biochemical experiments, and in individual cages for behavioural experiments. For SBFSEM experiments, all procedures were performed in full accordance with the recommendations of the European Community Council Directives (86/609/EEC), the French National Committee (87/848) and the requirements of the United Kingdom Animals (Scientific Procedures) Act 1986, AWERB Newcastle University (ID: 374).

### Antibodies

The primary antibodies used were: rabbit anti-dsRed (Living Colors® Clontech, used at 1/1000), rabbit anti-Vangl2 ^58^, homemade, used in IHC at 1/250 and in blots at 1/500), rabbit anti-synaptoporin (Synaptic Systems, used at 1/1000), rabbit anti-synapsin1 (Synaptic Systems, used at 1/1000), mouse anti-bassoon (Enzo Life Sciences, used at 1/3000), rabbit anti-GPC4 (Proteintech used at 1/2000 for blots and 1/200 for IHC), rabbit anti-GPR158 (Sigma-Aldrich used at 1/2000 for blots and 1/250 for IHC), chicken anti-GFP (Abcam, used at 1/15000), mouse anti-HA (BioLegend, used at 1/1000), mouse anti-myc (BioLegend used at 1/1000), mouse anti-PSD95 (ThermoFisher Scientific, used at 1/500), guinea pig anti-DCX (Millipore, used at 1/1000), mouse anti-ZNT3 (Synaptic System, used at 1/500), mouse anti-Tau (Millipore used at 1/1000), mouse anti-GAPDH (Millipore used at 1/5000). Alexa Fluor and other secondary antibodies were from ThermoFisher and Jackson ImmunoResearch and all were used at a dilution in between 1/1000 and 1/4000. Goat anti-mouse or anti-rabbit secondary antibodies coupled to horseradish peroxidase (HRP) (Jackson ImmunoResearch), donkey anti-guinea pig HRP (Jackson ImmunoResearch) and mouse anti-rabbit HRP (Jackson ImmunoResearch) were used in between 1/20000 and 1/40000 for Western blots experiments.

### Plasmids and viruses

The GFP-Vangl2 construct comes from ^58^. The HA-GRP158, HA-GPC4 constructs were kindly provided by Prof. de Wit (VIB-KU Leuven, Belgium) ^10^. The GPC4-myc construct was purchased from Origen. The adeno-associated viruses AAV2/9-CaMKII(0.4)-mCherry-WPRE and AAV2/9-CamKII(0.4)-mCherry-2A-mVangl2-WPRE were purchased from Vector Biolab ^13^.

### Cell culture, transfection, and immunocytochemistry

#### Hippocampal neurons

Neurons were cultured as described in ^55^. Neurons were maintained in a humidified incubator at 37°C with 5% CO_2_. Cells were transfected at DIV15 using calcium phosphate (CaP) precipitation as described in ^59^. For cDNA quantities, GFP-Vangl2 was used at 1.5 μg per well in combination with HA-GPC4, HA-GPR158 at 2µg per well.

#### COS-7 cells

COS-7 (ATCC-American Type Culture Collection) were cultured according to the manufacturer’s instructions and maintained in a humidified incubator at 37°C with 5% CO_2_. For transfection, we used CaP precipitation as described in ^59,60^. The different cDNAs were prepared with 12.5 μl of CaCl2 and H2O until a total volume of 100 μl. For cDNA quantities, GFP-Vangl2 was used at 3 μg for single transfection and at 1.5 μg in combination with other cDNAs. HA-GPC4, HA-GPR158 and GPC4-myc were used at 2µg, 2µg and 4µg respectively. The resulting mix was applied dropwise to 100 μl of 2× HBS incubated for 30 min at RT in the dark and applied to the cells. The plates were then placed back in the incubator for 5 hours. After that, three washes with serum-free DMEM were performed, and the cells were placed in the incubator for 10 min. Last, serum-free DMEM was replaced by complete medium. After 24h, the transfected cells were fixed and processed for immunocytochemistry. Briefly, COS-7 cells were fixed with 4% sucrose, 4% PFA in PBS pH 7.4 for 10 min, permeabilised and blocked with 0.3% Triton X-100, 5% bovine serum albumin (BSA), and incubated with mouse anti-HA, mouse anti-myc, and chicken anti-GFP antibodies for 1h at room temperature (RT). After extensive washing, fluorescently tagged goat anti-mouse and anti-chicken secondary antibodies were diluted in PBS and incubated for 30 min and coverslips were mounted using Fluoromount-G mounting medium (Electron Microscopy Sciences).

### Stereotaxic viral injection into neonatal and adult mice

Mouse pups (P5) were anaesthetised by inhalation of a 2% isoflurane/concentrated O_2_ mixture. 6-to 8-week-old mice were injected with a buprenorphine solution (0.15 mg/kg, intraperitoneally) 15 to 20 min before the start of the procedure. Prior to the procedure, adult mice were anaesthetised by inhalation of a 4% isoflurane/air mixture. Animals were considered anaesthetised when all movement and reflexes ceased.

Anaesthetised pups were secured in a digital stereotaxic frame with a neonatal mouse adaptor (World Precision Instruments, Inc., Sarasota, FL, USA) and maintained on 2% isoflurane/concentrated O_2_ inhalation. For adult mice, the injection site was shaved and disinfected, the eyes were protected with drops of an aqueous-based ocular preservative, and the animal was then mounted in a digital stereotaxic frame maintained on a 2% isoflurane/air inhalation mixture. Prior to surgical incision, 5 μl lidocaine for P5 pups (5 mg/kg) or 10 μl lidocaine (5 mg/kg) for adult mice was injected intradermally at the incision site for local anaesthesia.

Holes the size of the injection needle were made in the skull with a fine needle for P5 pups or with a microdrill for adult mice. Injections were distributed using a 33G for P5 pups and 34G Nanofil needle for adults attached to a Nanofil syringe (World Precision Instruments, Inc., Sarasota, FL, USA) coupled to a UMP3 UltraMicroPump (WPI). For Mf labelling and/or re-expression of *vangl2*, 100 nl (P5 pups) or 300 nl (6-to 8-week-old mice) of AAV2/9-CaMKII (0.4)-mCherry-WPRE virus (titre of 1. 9×10^12^ gcp/ml diluted in sterile PBS, Vector Biolabs) or AAV2/9-CaMKII (0.4)-mCherry-2A-mVangl2-WPRE virus (5.9×10^13^ gcp/ml or 1.7×10^13^ gcp/ml in sterile PBS, Vector Biolabs) were injected bilaterally into the dentate gyrus zone (P5 pups coordinates: Y: ∼1 mm from lambda; X: ±0.75 mm; Z: −1.65 mm), adult mice coordinates: X=±1.6, Y=-2, Z=-1.9 for mutants; X=±1.25, Y=-2, Z=-2 for controls) After surgery, P5 pups were allowed to recover on a warm heating pad and then returned to their mother until P25. After suturing, adult animals received a subcutaneous injection of warm 0.9% NaCl solution and were then allowed to recover in a clean individual cage placed in a heated chamber until full mobility was restored. After 48 h of post-operative observation, the mice were returned to the group housing.

### Enriched environement paradigm

For EE experiments, pairs of Emx1-Vangl2 and control male littermates were housed either in normal-sized shared cages without additional objects (6-7 mice per cage; control conditions) or in large (rat) cages equipped with 2 wheels per cage, toys, and hiding places for exploration (6 mice per cage; EE conditions). The objects in the EE conditions were moved every 2 days and changed completely every week. EE was started at 6 weeks of age, one week after surgery (see *Stereotaxic viral injection into neonatal and adult mice*) and continued for 5 weeks.

### Immunofluorescence on brain sections

Mice were terminally anaesthetised with intraperitoneal injections of pentobarbital and lidocaine (50 mg/kg body weight). For Vangl2 and mCherry, mice were injected with heparine and then transcardially perfused with phosphate buffer (PB) pH 7.4 followed by 4% PFA, brains were removed and postfixed in 4% PFA in PB for 2 to 24h at 4°C. 40-50 μm coronal vibratome sections (VT1000S vibratome, Leica) were processed for immunohistochemistry with anti-Vangl2 and anti-dsRed antibodies for 12 to 24h at 4°C. Alexa Fluor secondary antibodies were incubated for 1h at RT. Sections were mounted using ProLong Gold antifade medium (Life Technologies) or Fluoromount G medium.

For synaptic markers, brains were removed and frozen in embedding medium (M-1 Embedding Matrix, ThermoFisher) before storage at −80°C for at least 12h. 20-µm sections were fixed in either ice-cold 4% PFA in PBS or −20°C pure methanol for 5 min, permeabilised and incubated with 5% BSA, 5% NGS in PBS for 2h. Anti-GPR158, anti-GPC4, anti-synaptoporin, anti-synapsin1, and anti-bassoon antibodies were incubated for 12 to 24 h at 4°C. Alexa Fluor secondary antibodies were added for 1h at RT. Sections were mounted using ProLong Gold antifade medium.

For imaging we used a Zeiss AxioImager Z1 (63x/1.40 NA oil objective) equipped with an AxioCam MRm and the Zen software (Zeiss) and a LED light source Colibri 7 from Zeiss (wavelengths: UV 385/30 nm, V 423/44 nm, B 469/38 nm, C 511/44 nm, G 555/30 nm, Y 590/27 nm, R 631/33 nm), or a confocal Leica DM2500 TCS SPE laser scanning microscope with a 561 nm excitation wavelength and a 63x Leica oil objective. Images of the Mf termination zone in the CA3 *sl* region of the hippocampus were acquired at 300 nm intervals in the z-axis. Camera aperture, magnification, illumination and exposure time were fixed for all images. A minimum of 6 stacks were randomly acquired from each animal of each genotype and age. High-resolution 3D stacks were generated using Imaris 9.2.1 software (Bitplane, Oxford Instruments). The volume, surface area and number of filopodial processes per MfB were then determined for each 3D MfB.

### Serial block face scanning electron microscopy (SBFSEM), 3D reconstructions, and quantifications

P21 male mice were terminally anaesthetised by a short inhalation of isoflurane (0.05% in air), followed by an intramuscular injection of ketamine (100 mg/kg) and xylazine (10 mg/kg). Mice were then perfused intracardially with 5000 U heparin in PBS, followed by 200 ml of 2% PFA + 2.5% glutaraldehyde in PBS. Brains were dissected and stored at 4°C until shipped in 0.01% Na-azide in PBS at 4°C and on arrival were returned to 2% PFA + 2.5% glutaraldehyde in PBS at 4°C. The experimenters were blind to the genotype of the mice at all times. The code was not broken until all reconstructions, modelling, and analyses had been completed.

100-μm sections (Leica VT1000 S, Leica Biosystems) were trimmed to include the region of interest (ROI). Osmium impregnation began by rinsing samples in PBS, then 3% potassium ferrocyanide in 2% osmium tetroxide in PBS at 4°C for 1h. Samples were rinsed in ddH_2_0 at RT and incubated in freshly prepared and filtered 1% (w/v) thiocarbohydrazide for no more than 20 min to avoid crystallisation and precipitation. They were then rinsed in ddH_2_0, followed by 2% osmium tetroxide in ddH_2_O for 30 min, rinsed and stained *en bloc* in 1% uranyl acetate in ddH_2_0 overnight at 4°C. The next day, samples were rinsed in ddH_2_0, placed in Walton’s lead-aspartate solution at 60°C for 30 min, then rinsed in ddH_2_0 as before at RT. Dehydration through a graded ethanol series: 25%, then 10 min at 4°C each change (1 x 50%, 1 x 70%, 1 x 90%, 2 x 100%), followed by 3 times in 100% acetone for 30 min at RT. Samples were infiltrated by embedding in increasing concentrations of Durcupan resin (Sigma-Aldrich) mixed with acetone, starting with 50% resin and increasing to 70% after 1h and 90% after a further 1h at RT. Samples were then transferred to aluminium boats filled with 100% Durcupan resin for 4h at RT. The sections were then flat embedded and polymerised at 60°C for 48 h. Specimens containing the ROI were then mounted on metal stubs for SBFSEM ^7,61^. Samples were imaged on a Zeiss Sigma VP scanning electron microscope with Gatan 3View to collect images using 50 nm z-steps for control samples and 60 nm for Emx1-Vangl2 samples, measuring the same parameters as in ^7^.

3D models were generated from the SBFSEM image stacks using the publicly available Microscope Image Browser (MIB; http://mib.helsinki.fi/) software ^62^. The resulting 3D profiles were used to generate volumetric reconstructions and provide statistics on the volume of pre- and postsynaptic structures of interest. A total of 6 datasets were generated where postsynaptic dendrites were first traced and reconstructed in their entirety within the block. Spines, TEs, post-synaptic densities (PSDs), and presynaptic elements were then isolated and reconstructed individually in a similar manner. Spines and TEs were defined as a protrusion from the dendritic shaft, typically with a narrow neck. PSDs were defined as increased electron density and thickening of the postsynaptic membrane at an interface between presynaptic and postsynaptic structures. MfB were defined visually by their structure surrounding the previously isolated TE/spines. Reconstructions were constrained by the edges of the image block. The final 3D models and their morphological measurements were obtained at full resolution using Imaris 9.2.1 software (Bitplane).

### Western blot

For each set of experiments, 2 hippocampi from 3-week-old mice of each genotype were processed as previously described ^63^. Protein concentrations were measured using a BCA assay (Pierce, Rockford, IL), and equal amounts of proteins were separated by SDS-PAGE (10% or 4-20% gradient gels) and transferred to Immobilon-P membranes (Merck-Millipore). After chemiluminescence detection, the membranes were scanned using an imaging densitometer (BIORAD, Hercules, CA). Quantification was performed as previously described ^62^.

### Electrophysiological recordings

#### Slice preparation

21 to 32 postnatal days male mice were used. The methods for the slice preparation has been previously described in ^13^. In brief, animals were anaesthetised under isoflurane and decapitated. Brain was removed and prepared for slicing in an ice-cold cutting solution containing (in mM): 87 NaCl, 25 NaHCO_3_, 10 glucose, 75 sucrose, 2.5 KCl, 1.25 NaH_2_PO_4_, 0.5 CaCl_2_ and 7 MgSO_4,_ saturated with 95% O_2_ and 5% CO_2_. Then, 300 µm parasagittal hippocampal slices were cut on a vibratome in ice-cold and oxygenated cutting solution. After sectioning, slices were allowed to recover for 30 min to 1 h at 33°C and then kept at RT for the duration of the experiment. Slices were transferred to a recording chamber continuously superfused at a rate of 3-4 ml/min with an oxygenated artificial CSF at RT (aCSF; 95% O_2_ and 5% CO_2_) containing (in mM): 125 NaCl, 2.5 KCl, 2.5 CaCl_2_, 1.3 MgSO_4_, 1.25 NaH_2_PO_4_, 25 NaHCO_3_, 10 glucose, pH 7.4. Voltage clamp recordings from CA3 PCs were made using borosilicate pipettes (4-8 MΩ) filled with methane-sulfonate solution as previously described in ^13^.

#### Evoked excitatory postsynaptic currents (eEPSCs) recordings

eEPSCs were recorded in CA3 PCs in voltage-clamp mode by stimulating the DG hilus with a glass pipette to activate Mfs. Mf CA3 PC eEPSCs were evoked by minimal intensity stimulation according to the previously described protocol ^64,65^.

#### AMPAR/NMDAR ratio recordings

Cells were clamped at −70 mV to isolate AMPAR/KAR response in the presence of the NMDA receptor antagonist D-APV (50 μM) added to ACSF. After 15 min of D-APV washing, the AMPAR/KAR antagonist NBQX (20 µM) was added to record NMDAR EPSC amplitudes at +40 mV. For each pharmacological condition, 30 sweeps were averaged.

#### Long-term potentiation recordings

LTP was induced with HFS consisting of 100 Hz stimulation trains lasting 1 s and repeated three times at 10 s intervals. Following the HFS, eEPSCs were recorded for 50 min at 0.1 Hz. The magnitude of LTP was calculated the last 10 min of the recording.

#### Drugs

eEPSC and LTP recordings were performed in aCSF with the GABA_A_ receptor antagonist bicuculline methiodide (20 μM) and the NMDA receptor antagonist D-APV (50 μM)^65^. The metabotropic glutamate receptor 2/3 agonist (2S,1′R,2′R,3′R)-2-(2,3-dicarboxycyclopropyl)glycine (DCG-IV; 1 μM) was applied at the end of some experiments to selectively block Mf responses ^10^. All drugs were obtained from Tocris Cookson (Bristol, UK), SigmaAldrich (St. Louis, MO), HelloBio (Bristol, UK).

#### Data acquisition

Recordings were made *via* Patchmaster 2.71 using an EPC10 amplifier (HEKA Elektronik, Lambrecht/ Pfalz, Germany), filtered at 0.5–1 kHz, digitised at 5-10 kHz. Analysis was performed using Neuromatic3.0 (www.neuromatic.thinkrandom.com) written within the Igor Pro 6.37 environment (WaveMetrics, Lake Oswego, OR).

### Behavioural tests

Experiments were performed with control and Emx1-Vangl2 littermates. All animals were male mice, 10-12 weeks of age at the start of the behavioural tests. Mice were kept in collective housings in standard conditions (food and water ad libitum; 22±2°C, 45-65% hydrometry, 12-12 h light/dark cycle, light on at 7:00; experiments performed between 9:00 and 17:00) except for the radial maze experiments where they were kept in individual housings. The most stressful tasks were performed last to minimise interference.

#### Locomotor activity

Locomotor activity was assessed in photocell-based activity chambers under light-dark environmental conditions using interconnected individual Plexiglas chambers equipped with infrared sensors, as previously published ^63^. Rearing and horizontal activity data were collected for each mouse over a 3-h time course (response to novelty in 10-min bouts).

#### Elevated plus-maze

The Plexiglas plus maze apparatus (Noldus) had two open and two closed arms radiating outwards from a central open square and was 60 cm above the floor. The maze was illuminated by halogen light places in the 4 corners of the room, set to 30 lux. The mice were placed in the centre of the maze with free access to all arms for 5 min. A video tracking system (Ethovision version XT17, Noldus Technology, Wageningen, The Netherlands) was used to analyse the percentage of time spent and the number of entries into the open arms.

#### Dark/light exploration assay

Dark/light was conducted in a two-chamber shuttle box with an opaque partition in the middle. The dark chamber was made of black Plexiglas walls, while the light chamber was white and illuminated by LED light (around 200 lux). The mice were first placed in the dark chamber and allowed to move freely between the chambers for 5 min. The percentage of time spent and the number of entries into the light chamber were analysed using a video tracking system (Ethovision).

#### Spontaneous alternation in the Y-maze

Spontaneous alternation was assessed in a white plastic Y-maze with 3 arms of similar appearance, separated by 120°, and illuminated by the same light. Mice were placed at the end of one of the arms and allowed to explore the maze for 5 min. Start arm allocation was counterbalanced within experimental groups. A correct alternation was defined as a non-repeated entry into the arms for three consecutive entries.

#### Spatial learning and memory in the Morris water maze

This test has been previously described in ^63^. In brief, spatial memory was tested in a circular pool filled with opaque water kept at 19-20°C. Mice were trained to swim to a platform to escape from the water for 1 min. Data were collected using a video tracking camera (Viewpoint, Lyon, France). To measure performance in the hidden trials, escape latency, distance to the target and swimming speed were analysed as outcome measures for each session. If the mice found the platform, they were allowed to stay on it for 15 s. If they did not, they were guided to the platform where they were allowed to stay for 15 s. Mice were released from a different starting point on each trial, and different sequences of starting points were used each day. *Visual training*: During 5 consecutive sessions, mice were trained to swim to an unsubmerged platform indicated by a ball suspended 7.5 cm above the platform. *Spatial training*: For 10 days, mice were trained to swim to a submerged platform (1.5 cm below the water surface) using extramaze spatial cues placed in the room. *Probe trials*: The platform was removed from the pool 24 h after the last hidden platform training trial. The time the mice spent swimming in the target quadrant were measured over a 60 s trial.

#### Relational/Declarative memory in the radial maze

One week before and until the end of the experiments, the mice were subjected to partial food deprivation (82-85% of their free food weight) and handled daily. We used three identical fully automated 8-arm radial maze made of grey Plexiglas. Only extra-maze cues were available. *Habituation*: Each mouse underwent two days of habituation, in which all arms were baited and open, and had to visit each arm and eat all pellets to complete the session. *Training*: Each mouse was assigned three pairs of adjacent arms (pairs A, B, and C) for the entire training period. One arm of each pair is always baited. Within each daily training session, the mouse was exposed to 20 consecutive presentations (trials) of the three pairs in a pseudo-random order, separated by an inter trial interval (ITI) of 20 s (Figure 6A). A trial was considered successful if the mouse chose the baited arm. Training lasted for a minimum of 5 days and ended when a mouse achieved an average performance of 72.5% for all three pairs on two days. Mice that did not complete each daily session in a maximum of one hour and mice that did not complete training in 12 days were excluded from the analysis (here 3 control and 3 Emx1-Vangl2 mice were excluded). *Test*: When a mouse completed training, it was subjected to a test session of 20 trials. During this session, each mouse was assigned three pairs of arms: the known pair C (positive control pair), a new pair N (negative control pair), and a recombination pair AB using the unbaited arm of pair A and the baited arm of pair B. The percentage of performance obtained for this pair AB served as an index of memory flexibility.

#### Every day-like memory in the radial maze

This task has been described in detail in ^25^. The apparatus, food deprivation, handling, and habituation phase were the same used as in the previous behavioural test. *Habituation*: Each mouse underwent one day of habituation as described above. *Training*: Each mouse was assigned three pairs of adjacent arms (pairs A, B, and C) for the entire training session. Within each daily training session, the mouse was exposed to 23 consecutive presentations (trials) of the three pairs in a pseudo-random order, separated by a 10 s ITI. For the first presentation of each pair, both arms were baited. Then, for subsequent presentations, the reward was in the arm not visited during the previous trial with the same pair, so that the mouse had to alternate between the arms for each pair. The difficulty of the task was balanced over three days of training, so we compared performance between blocks of three days. Mice that did not complete each daily session in a maximum of one hour were excluded from the analysis.

#### Contextual fear conditioning

The contextual fear conditioning used is based on a tone-shock unpairing procedure, where the tone does not predict the shock (pseudorandomly distribution of the tone and shock) making the background context the only predictor ^66–68^. We used two different chambers for conditioning. The conditioning chamber is the same as used in ^68^. The neutral chamber was made of opaque grey plastic (30×24×22 cm) with a plastic floor. On the first day, mice were placed in the neutral chamber for a 3-min habituation period. 24 h later, mice were placed in the (translucent) conditioning chamber for 4 min: 100 s after being placed in the chamber, animals received a footshock (0.4 mA); after a 20 s delay, a tone was presented (65 dB, 1 kHz, 15 s); after a 30 s delay, the same tone was presented followed by the same footshock 30 s after; finally, after 20 s the mice were replaced in their home cage. 24 h later, mice were subjected to 4 tests and freezing behaviour was recorded as an index of fear. First, mice were placed in the neutral chamber for 6 min and the tone (65 dB, 1 kHz, 2 min) was presented after a 2min delay. 2 h later, the mice were placed in the conditioning chamber for 6 min, without tone or shock. During the first 3 min, the electrical grid was hidden under a plastic plate to prevent its use as a cue. This plastic plate was removed at the end of the test. 2 h later, the mice were placed in the neutral chamber for 6 min and a similar, but different, tone (65 dB, 2 kHz, 2 min) was presented after a 2 min delay in order to assess any generalisation of fear. With the same aim, 2 h later, mice were placed in the neutral chamber for 6 min and a very different noise (i.e. white noise, 65 dB) was presented for 2 min after a 2 min delay.

### Statistical analysis

All data are presented as mean ± SEM, and p ≤ 0.05 was considered statistically significant (*p<0.05, **p<0.01, ***p<0.001, p<0,0001). When necessary, the assumption of normality was tested. All calculations were performed using GraphPad Prism version 8 for Windows. For 3D reconstruction experiments, normality was tested using the Shapiro-Wilk test. If the normality test failed, we used the Mann-Whitney rank sum test to assess statistical significance between groups.

Electrophysiological datasets were tested for normality using D’Agostino and Pearson test. If datasets passed the test, they were analysed using Student’s unpaired t-test with Welch’s correction. Otherwise, the datasets were analysed using nonparametric unpaired t-tests (Mann–Whitney). For paired-pulse ratio experiments, comparisons between two genotypes were performed using two-way ANOVA test (Bonferroni’s multiple comparisons test). N mentioned in the paper represents the number of mice used in an experiment; n mentioned in the paper represents a single measurement from a single cell or a measurement of a single structure. No statistical methods were used to predetermine sample sizes, but our sample sizes are similar to those generally employed in the field.

For behavioural tests, one-way ANOVA was used to analyse the effect of genotype on time spent in each quadrant of the water maze in the probe test. Two-way ANOVA was used to analyse the effect of training days and genotype on declarative and working memory performance in the RM test. The Bonferroni’s post hoc test was used when appropriate. Student’s t-test was used to compare control and Emx1-Vangl2 mice in the plus-maze, dark/light test, locomotor rearing, Welch t-test was used for Y-maze tests and Mann-Whitney test was used for declarative memory assessment.

## Acknowledgments

We thank: Prof. D. Henderson for the Vangl2 flox mice and Dr J. De Wit for the GPC4 and GPR58 cDNAs. We thank the personnel of the Animal Facility of the Neurocentre Magendie for the mouse care (notably M. Jacquet, R. Racunica and J-B Bernard) and of the PUMA genotyping facility (notably D. Gonzales and her team). Confocal imaging was done at the Bordeaux Imaging Center a service unit of the CNRS-INSERM-Bordeaux University, a member of the national infrastructure France BioImaging (ANR-10-INBS-04). All these facilities were funded by the Labex B.R.A.I.N. (ANR-10-LABX-43). We also thank the Electron Microscopy Research Services, Faculty of Medical Sciences, Newcastle University. We also would like to thank the members of the Montcouquiol/Sans laboratory for critical reading and comments on the manuscript.

This work was supported by INSERM, ANR MossyPCP ANR-12-BSV4-0016-01 (NS, CM, AM), ANR THORNEX ANR-20-CE16-0028-01 (NS, AM), La Fondation pour la Recherche Médicale “Equipe FRM 2016” DEQ20160334899 (MM). ND was an MESRI Ph.D.fellow and received a 4^th^ year fellowship from the Labex B.R.A.I.N. (ANR-10-LABX-43).

## Author contributions

Conceptualisation, N.D., N.S., M.M.

Methodology, N.D., M.G., M.M.M., C.M., C.R., A.D., A.M., M.M., and N.S

Investigation, N.D., M.G., M.M.M., A.Q.

Formal Analysis, N.D, M.G., M.M.M., A.Q., A.M.

Supervision, M.M.M., G.B., C.R., A.M., M.M., and N.S.;

Resources, D.J.H.

Writing – Original Draft, N.D., M.M., and N.S.;

Writing – Review & Editing, All authors discussed the results and edited the manuscript; Project Administration, N.S.; Funding Acquisition, N.S., M.M., A.M., and C.M.

## Data and materials availability

All data needed to evaluate the conclusions in the paper are present in the paper and/or the Supplementary Materials. All reagents are available upon request. The mouse line can be provided pending a completed material transfer agreement. Requests for all materials should be submitted to N.S. at.

## References

1. Lanore, F., Labrousse, V.F., Szabo, Z., Normand, E., Blanchet, C., and Mulle, C. (2012). Deficits in Morphofunctional Maturation of Hippocampal Mossy Fiber Synapses in a Mouse Model of Intellectual Disability. J. Neurosci. 32, 17882–17893. 10.1523/JNEUROSCI.2049-12.2012.

2. Witton, J., Padmashri, R., Zinyuk, L.E., Popov, V.I., Kraev, I., Line, S.J., Jensen, T.P., Tedoldi, A., Cummings, D.M., Tybulewicz, V.L.J., et al. (2015). Hippocampal circuit dysfunction in the Tc1 mouse model of Down syndrome. Nat Neurosci 18, 1291–1298. 10.1038/nn.4072.

3. Wilke, S.A., Raam, T., Antonios, J.K., Bushong, E.A., Koo, E.H., Ellisman, M.H., and Ghosh, A. (2014). Specific disruption of hippocampal mossy fiber synapses in a mouse model of familial Alzheimer’s disease. PLoS ONE 9, e84349. 10.1371/journal.pone.0084349.

4. Silva, S.V. da, Zhang, P., Haberl, M.G., Labrousse, V., Grosjean, N., Blanchet, C., Frick, A., and Mulle, C. (2019). Hippocampal Mossy Fibers Synapses in CA3 Pyramidal Cells Are Altered at an Early Stage in a Mouse Model of Alzheimer’s Disease. J. Neurosci. 39, 4193– 4205. 10.1523/JNEUROSCI.2868-18.2019.

5. Amaral, D.G., and Dent, J.A. (1981). Development of the mossy fibers of the dentate gyrus: I. A light and electron microscopic study of the mossy fibers and their expansions. Journal of Comparative Neurology 195, 51–86. 10.1002/cne.901950106.

6. Urban, N.N., Henze, D.A., and Barrionuevo, G. (2001). Revisiting the role of the hippocampal mossy fiber synapse. Hippocampus 11, 408–417. 10.1002/hipo.1055.

7. Wilke, S.A., Antonios, J.K., Bushong, E.A., Badkoobehi, A., Malek, E., Hwang, M., Terada, M., Ellisman, M.H., and Ghosh, A. (2013). Deconstructing complexity: serial block-face electron microscopic analysis of the hippocampal mossy fiber synapse. J. Neurosci. 33, 507–522. 10.1523/JNEUROSCI.1600-12.2013.

8. Zhang, P., Lu, H., Peixoto, R.T., Pines, M.K., Ge, Y., Oku, S., Siddiqui, T.J., Xie, Y., Wu, W., Archer-Hartmann, S., et al. (2018). Heparan Sulfate Organizes Neuronal Synapses through Neurexin Partnerships. Cell 174, 1450–1464.e23. 10.1016/j.cell.2018.07.002.

9. Kamimura, K., and Maeda, N. (2021). Glypicans and Heparan Sulfate in Synaptic Development, Neural Plasticity, and Neurological Disorders. Front Neural Circuits 15. 10.3389/fncir.2021.595596.

10. Condomitti, G., Wierda, K.D., Schroeder, A., Rubio, S.E., Vennekens, K.M., Orlandi, C., Martemyanov, K.A., Gounko, N.V., Savas, J.N., and de Wit, J. (2018). An Input-Specific Orphan Receptor GPR158-HSPG Interaction Organizes Hippocampal Mossy Fiber-CA3 Synapses. Neuron 100, 201–215.e9. 10.1016/j.neuron.2018.08.038.

11. Ma, K.-G., Hu, H.-B., Zhou, J.-S., Ji, C., Yan, Q.-S., Peng, S.-M., Ren, L.-D., Yang, B.-N., Xiao, X.-L., Ma, Y.-B., et al. (2022). Neuronal Glypican4 promotes mossy fiber sprouting through the mTOR pathway after pilocarpine-induced status epilepticus in mice. Experimental Neurology 347, 113918. 10.1016/j.expneurol.2021.113918.

12. Davis, G.W. (2000). The Making of a Synapse: Target-Derived Signals and Presynaptic Differentiation. Neuron 26, 551–554. 10.1016/S0896-6273(00)81190-3.

13. Robert, B.J.A., Moreau, M.M., Dos Santos Carvalho, S., Barthet, G., Racca, C., Bhouri, M., Quiedeville, A., Garret, M., Atchama, B., Al Abed, A.S., et al. (2020). Vangl2 in the Dentate Network Modulates Pattern Separation and Pattern Completion. Cell Reports 31, 107743. 10.1016/j.celrep.2020.107743.

14. Nagaoka, T., and Kishi, M. (2016). The planar cell polarity protein Vangl2 is involved in postsynaptic compartmentalization. Neuroscience Letters 612, 251–255. 10.1016/j.neulet.2015.12.009.

15. Dos-Santos Carvalho, S., Moreau, M.M., Hien, Y.E., Garcia, M., Aubailly, N., Henderson, D.J., Studer, V., Sans, N., Thoumine, O., and Montcouquiol, M. (2020). Vangl2 acts at the interface between actin and N-cadherin to modulate mammalian neuronal outgrowth. eLife 9, e51822. 10.7554/eLife.51822.

16. Henze, D.A., Urban, N.N., and Barrionuevo, G. (2000). The multifarious hippocampal mossy fiber pathway: a review. Neuroscience 98, 407–427. 10.1016/S0306-4522(00)00146-9.

17. Love, A.M., Prince, D.J., and Jessen, J.R. (2018). Vangl2-dependent regulation of membrane protrusions and directed migration requires a fibronectin extracellular matrix. Development 145. 10.1242/dev.165472.

18. Balaraju, A.K., Hu, B., Rodriguez, J.J., Murry, M., and Lin, F. (2021). Glypican 4 regulates planar cell polarity of endoderm cells by controlling the localization of Cadherin 2. Development 148. 10.1242/dev.199421.

19. Galimberti, I., Gogolla, N., Alberi, S., Santos, A.F., Muller, D., and Caroni, P. (2006). Long-Term Rearrangements of Hippocampal Mossy Fiber Terminal Connectivity in the Adult Regulated by Experience. Neuron 50, 749–763. 10.1016/j.neuron.2006.04.026.

20. Gogolla, N., Galimberti, I., Deguchi, Y., and Caroni, P. (2009). Wnt Signaling Mediates Experience-Related Regulation of Synapse Numbers and Mossy Fiber Connectivities in the Adult Hippocampus. Neuron 62, 510–525. 10.1016/j.neuron.2009.04.022.

21. Henze, D.A., McMahon, D.B.T., Harris, K.M., and Barrionuevo, G. (2002). Giant Miniature EPSCs at the Hippocampal Mossy Fiber to CA3 Pyramidal Cell Synapse Are Monoquantal. Journal of Neurophysiology 87, 15–29. 10.1152/jn.00394.2001.

22. Apóstolo, N., Smukowski, S.N., Vanderlinden, J., Condomitti, G., Rybakin, V., ten Bos, J., Trobiani, L., Portegies, S., Vennekens, K.M., Gounko, N.V., et al. (2020). Synapse type-specific proteomic dissection identifies IgSF8 as a hippocampal CA3 microcircuit organizer. Nat Commun 11, 5171. 10.1038/s41467-020-18956-x.

23. Marighetto, A., Etchamendy, N., Touzani, K., Torrea, C.C., Yee, B.K., Rawlins, J.N., and Jaffard, R. (1999). Knowing which and knowing what: a potential mouse model for age-related human declarative memory decline. Eur J Neurosci 11, 3312–3322. 10.1046/j.1460-9568.1999.00741.x.

24. Etchamendy, N., Enderlin, V., Marighetto, A., Pallet, V., Higueret, P., and Jaffard, R. (2003). Vitamin A deficiency and relational memory deficit in adult mice: relationships with changes in brain retinoid signalling. Behav Brain Res 145, 37–49. 10.1016/s0166-4328(03)00099-8.

25. Mingaud, F., Mormede, C., Etchamendy, N., Mons, N., Niedergang, B., Wietrzych, M., Pallet, V., Jaffard, R., Krezel, W., Higueret, P., et al. (2008). Retinoid Hyposignaling Contributes to Aging-Related Decline in Hippocampal Function in Short-Term/Working Memory Organization and Long-Term Declarative Memory Encoding in Mice. J Neurosci 28, 279–291. 10.1523/JNEUROSCI.4065-07.2008.

26. Al Abed, A.S., Sellami, A., Brayda-Bruno, L., Lamothe, V., Noguès, X., Potier, M., Bennetau-Pelissero, C., and Marighetto, A. (2016). Estradiol enhances retention but not organization of hippocampus-dependent memory in intact male mice. Psychoneuroendocrinology 69, 77–89. 10.1016/j.psyneuen.2016.03.014.

27. Nagaoka, T., Ohashi, R., Inutsuka, A., Sakai, S., Fujisawa, N., Yokoyama, M., Huang, Y.H., Igarashi, M., and Kishi, M. (2014). The Wnt/Planar Cell Polarity Pathway Component Vangl2 Induces Synapse Formation through Direct Control of N-Cadherin. Cell Reports 6, 916–927. 10.1016/j.celrep.2014.01.044.

28. Thakar, S., Wang, L., Yu, T., Ye, M., Onishi, K., Scott, J., Qi, J., Fernandes, C., Han, X., Yates, J.R., et al. (2017). Evidence for opposing roles of Celsr3 and Vangl2 in glutamatergic synapse formation. Proc Natl Acad Sci U S A 114, E610–E618. 10.1073/pnas.1612062114.

29. Montcouquiol, M., and Kelley, M.W. (2020). Development and Patterning of the Cochlea: From Convergent Extension to Planar Polarity. Cold Spring Harb Perspect Med 10, a033266. 10.1101/cshperspect.a033266.

30. Roszko, I., Sawada, A., and Solnica-Krezel, L. (2009). Regulation of convergence and extension movements during vertebrate gastrulation by the Wnt/PCP pathway. Semin Cell Dev Biol 20, 986–997. 10.1016/j.semcdb.2009.09.004.

31. Wang, M., de Marco, P., Capra, V., and Kibar, Z. (2019). Update on the Role of the Non-Canonical Wnt/Planar Cell Polarity Pathway in Neural Tube Defects. Cells 8, 1198. 10.3390/cells8101198.

32. Roszko, I., S. Sepich, D., Jessen, J.R., Chandrasekhar, A., and Solnica-Krezel, L. (2015). A dynamic intracellular distribution of Vangl2 accompanies cell polarization during zebrafish gastrulation. Development 142, 2508–2520. 10.1242/dev.119032.

33. Marlow, F., Zwartkruis, F., Malicki, J., Neuhauss, S.C., Abbas, L., Weaver, M., Driever, W., and Solnica-Krezel, L. (1998). Functional interactions of genes mediating convergent extension, knypek and trilobite, during the partitioning of the eye primordium in zebrafish. Dev Biol 203, 382–399. 10.1006/dbio.1998.9032.

34. Nychyk, O., Galea, G.L., Molè, M., Savery, D., Greene, N.D.E., Stanier, P., and Copp, A.J. (2022). Vangl2–environment interaction causes severe neural tube defects, without abnormal neuroepithelial convergent extension. Dis Model Mech 15, dmm049194. 10.1242/dmm.049194.

35. Galli, A., Roure, A., Zeller, R., and Dono, R. (2003). Glypican 4 modulates FGF signalling and regulates dorsoventral forebrain patterning in Xenopus embryos. Development 130, 4919–4929. 10.1242/dev.00706.

36. Bekirov, I.H., Nagy, V., Svoronos, A., Huntley, G.W., and Benson, D.L. (2008). Cadherin-8 and N-cadherin Differentially Regulate Pre-and Postsynaptic Development of the Hippocampal Mossy Fiber Pathway. Hippocampus 18, 349–363. 10.1002/hipo.20395.

37. van Stegen, B., Dagar, S., and Gottmann, K. (2017). Release activity-dependent control of vesicle endocytosis by the synaptic adhesion molecule N-cadherin. Sci Rep 7, 40865. 10.1038/srep40865.

38. Dagar, S., Teng, Z., and Gottmann, K. (2021). Transsynaptic N-Cadherin Adhesion Complexes Control Presynaptic Vesicle and Bulk Endocytosis at Physiological Temperature. Frontiers in Cellular Neuroscience 15.

39. Mirza, F.J., and Zahid, S. (2017). The Role of Synapsins in Neurological Disorders. Neurosci Bull 34, 349–358. 10.1007/s12264-017-0201-7.

40. Knaus, P., Marquèze-Pouey, B., Scherer, H., and Betzt, H. (1990). Synaptoporin, a novel putative channel protein of synaptic vesicles. Neuron 5, 453–462. 10.1016/0896-6273(90)90084-S.

41. Sara, Y., Bal, M., Adachi, M., Monteggia, L.M., and Kavalali, E.T. (2011). Use-Dependent AMPA Receptor Block Reveals Segregation of Spontaneous and Evoked Glutamatergic Neurotransmission. J Neurosci 31, 5378–5382. 10.1523/JNEUROSCI.5234-10.2011.

42. Sara, Y., Virmani, T., Deák, F., Liu, X., and Kavalali, E.T. (2005). An Isolated Pool of Vesicles Recycles at Rest and Drives Spontaneous Neurotransmission. Neuron 45, 563–573. 10.1016/j.neuron.2004.12.056.

43. Kavalali, E.T. (2015). The mechanisms and functions of spontaneous neurotransmitter release. Nat Rev Neurosci 16, 5–16. 10.1038/nrn3875.

44. Copley, C.O., Duncan, J.S., Liu, C., Cheng, H., and Deans, M.R. (2013). Postnatal Refinement of Auditory Hair Cell Planar Polarity Deficits Occurs in the Absence of Vangl2. J. Neurosci. 33, 14001–14016. 10.1523/JNEUROSCI.1307-13.2013.

45. Nicoll, R.A., and Schmitz, D. (2005). Synaptic plasticity at hippocampal mossy fibre synapses. Nature Reviews Neuroscience 6, 863–876. 10.1038/nrn1786.

46. Maruo, T., Mandai, K., Takai, Y., and Mori, M. (2016). Activity-dependent alteration of the morphology of a hippocampal giant synapse. Molecular and Cellular Neuroscience 71, 25–33. 10.1016/j.mcn.2015.12.005.

47. Rolls, E.T. (2016). Pattern separation, completion, and categorisation in the hippocampus and neocortex. Neurobiology of Learning and Memory 129, 4–28. 10.1016/j.nlm.2015.07.008.

48. Burghardt, N.S., Park, E.H., Hen, R., and Fenton, A.A. (2012). Adult-born hippocampal neurons promote cognitive flexibility in mice. Hippocampus 22, 1795–1808. 10.1002/hipo.22013.

49. Xavier, G.F., Oliveira-Filho, F.J., and Santos, A.M. (1999). Dentate gyrus-selective colchicine lesion and disruption of performance in spatial tasks: difficulties in “place strategy” because of a lack of flexibility in the use of environmental cues? Hippocampus 9, 668–681. 10.1002/(SICI)1098-1063(1999)9:6<668::AID-HIPO8>3.0.CO;2-9.

50. Lee, C.-H., and Lee, I. (2020). Impairment of Pattern Separation of Ambiguous Scenes by Single Units in the CA3 in the Absence of the Dentate Gyrus. J. Neurosci. 40, 3576–3590. 10.1523/JNEUROSCI.2596-19.2020.

51. Hughes, R.N. (2004). The value of spontaneous alternation behavior (SAB) as a test of retention in pharmacological investigations of memory. Neuroscience & Biobehavioral Reviews 28, 497–505. 10.1016/j.neubiorev.2004.06.006.

52. McLamb, R.L., Mundy, W.R., and Tilson, H.A. (1988). Intradentate colchicine disrupts the acquisition and performance of a working memory task in the radial arm maze. Neurotoxicology 9, 521–528.

53. Kesner, R. (2013). A process analysis of the CA3 subregion of the hippocampus. Frontiers in Cellular Neuroscience 7.

54. Hunsaker, M.R., Tran, G.T., and Kesner, R.P. (2009). A behavioral analysis of the role of CA3 and CA1 subcortical efferents during classical fear conditioning. Behav Neurosci 123, 624–630. 10.1037/a0015455.

55. Dorrego-Rivas, A., Ezan, J., Moreau, M.M., Poirault-Chassac, S., Aubailly, N., De Neve, J., Blanchard, C., Castets, F., Fréal, A., Battefeld, A., et al. (2022). The core PCP protein Prickle2 regulates axon number and AIS maturation by binding to AnkG and modulating microtubule bundling. Sci Adv 8, eabo6333. 10.1126/sciadv.abo6333.

56. Ramsbottom, S.A., Sharma, V., Rhee, H.J., Eley, L., Phillips, H.M., Rigby, H.F., Dean, C., Chaudhry, B., and Henderson, D.J. (2014). Vangl2-Regulated Polarisation of Second Heart Field-Derived Cells Is Required for Outflow Tract Lengthening during Cardiac Development. PLoS Genet 10, e1004871. 10.1371/journal.pgen.1004871.

57. Gorski, J.A., Talley, T., Qiu, M., Puelles, L., Rubenstein, J.L.R., and Jones, K.R. (2002). Cortical Excitatory Neurons and Glia, But Not GABAergic Neurons, Are Produced in the Emx1-Expressing Lineage. J Neurosci 22, 6309–6314. 10.1523/JNEUROSCI.22-15-06309.2002.

58. Montcouquiol, M., Sans, N., Huss, D., Kach, J., Dickman, J.D., Forge, A., Rachel, R.A., Copeland, N.G., Jenkins, N.A., Bogani, D., et al. (2006). Asymmetric Localization of Vangl2 and Fz3 Indicate Novel Mechanisms for Planar Cell Polarity in Mammals. J Neurosci 26, 5265– 5275. 10.1523/JNEUROSCI.4680-05.2006.

59. Sans, N., Wang, P.Y., Du, Q., Petralia, R.S., Wang, Y.-X., Nakka, S., Blumer, J.B., Macara, I.G., and Wenthold, R.J. (2005). mPins modulates PSD-95 and SAP102 trafficking and influences NMDA receptor surface expression. Nat Cell Biol 7, 1179–1190. 10.1038/ncb1325.

60. Piguel, N.H., Fievre, S., Blanc, J.-M., Carta, M., Moreau, M.M., Moutin, E., Pinheiro, V.L., Medina, C., Ezan, J., Lasvaux, L., et al. (2014). Scribble1/AP2 Complex Coordinates NMDA Receptor Endocytic Recycling. Cell Reports 9, 712–727. 10.1016/j.celrep.2014.09.017.

61. Tapia, J.C., Kasthuri, N., Hayworth, K., Schalek, R., Lichtman, J.W., Smith, S.J., and Buchanan, J. (2012). High contrast en bloc staining of neuronal tissue for field emission scanning electron microscopy. Nat Protoc 7, 193–206. 10.1038/nprot.2011.439.

62. Belevich, I., Joensuu, M., Kumar, D., Vihinen, H., and Jokitalo, E. (2016). Microscopy Image Browser: A Platform for Segmentation and Analysis of Multidimensional Datasets. PLOS Biology 14, e1002340. 10.1371/journal.pbio.1002340.

63. Moreau, M.M., Piguel, N., Papouin, T., Koehl, M., Durand, C.M., Rubio, M.E., Loll, F., Richard, E.M., Mazzocco, C., Racca, C., et al. (2010). The Planar Polarity Protein Scribble1 Is Essential for Neuronal Plasticity and Brain Function. J Neurosci 30, 9738–9752. 10.1523/JNEUROSCI.6007-09.2010.

64. Marchal, C., and Mulle, C. (2004). Postnatal maturation of mossy fibre excitatory transmission in mouse CA3 pyramidal cells: a potential role for kainate receptors. The Journal of Physiology 561, 27–37. 10.1113/jphysiol.2004.069922.

65. Sachidhanandam, S., Blanchet, C., Jeantet, Y., Cho, Y.H., and Mulle, C. (2009). Kainate Receptors Act as Conditional Amplifiers of Spike Transmission at Hippocampal Mossy Fiber Synapses. J Neurosci 29, 5000–5008. 10.1523/JNEUROSCI.5807-08.2009.

66. Desmedt, A., Garcia, R., and Jaffard, R. (1998). Differential Modulation of Changes in Hippocampal–Septal Synaptic Excitability by the Amygdala as a Function of Either Elemental or Contextual Fear Conditioning in Mice. J Neurosci 18, 480–487. 10.1523/JNEUROSCI.18-01-00480.1998.

67. Calandreau, L., Jaffard, R., and Desmedt, A. (2007). Dissociated roles for the lateral and medial septum in elemental and contextual fear conditioning. Learn Mem 14, 422–429. 10.1101/lm.531407.

68. Calandreau, L., Trifilieff, P., Mons, N., Costes, L., Marien, M., Marighetto, A., Micheau, J., Jaffard, R., and Desmedt, A. (2006). Extracellular Hippocampal Acetylcholine Level Controls Amygdala Function and Promotes Adaptive Conditioned Emotional Response. J Neurosci 26, 13556–13566. 10.1523/JNEUROSCI.3713-06.2006.

