## Supplemental figures for "The correct temporal connectivity of the DG-CA3 circuits involved in declarative memory processes depends on Vangl2-dependent planar cell polarity signaling"

[1]Univ. Bordeaux, Inserm, Neurocentre Magendie, U1215, F-33000 Bordeaux, France.
[2]Univ. Bordeaux, CNRS, IINS, UMR 5297, F-33000 Bordeaux, France.
[3]Biosciences Institute, The Medical School, Newcastle University, Newcastle upon Tyne, United Kingdom

### These authors contributed equally to this work;
* *Equal Senior contribution;*
✝ Late
Noémie Depret

**This pdf file includes :**

Supplementary Fig. 1

Supplementary Fig. 2 related to Fig. 2

Supplementary Fig. 3

Supplementary Fig. 4 related to Fig. 7

Supplementary Fig. 5

Supplementary Table 1

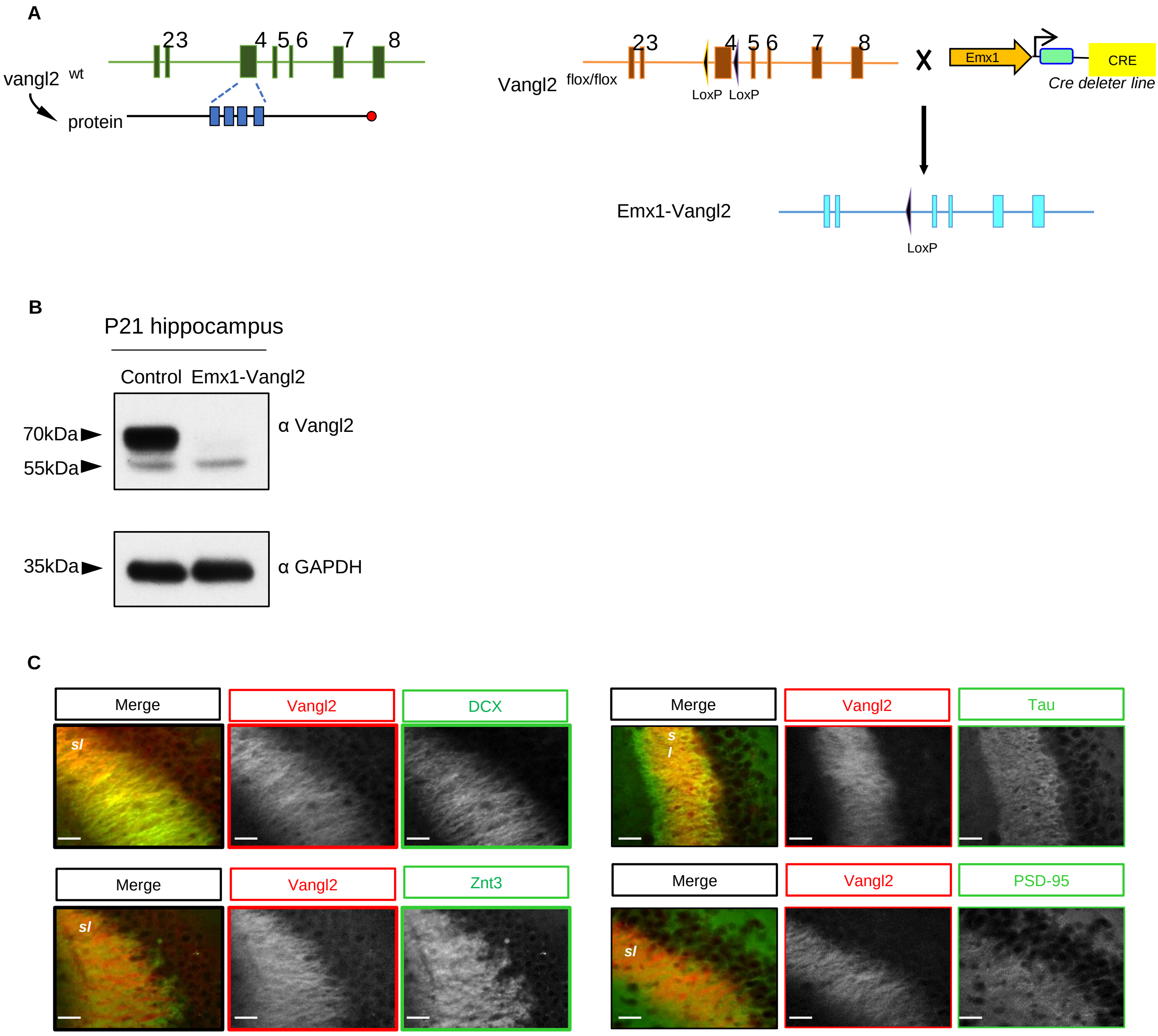


**Supplementary Fig. 1: Conditional *vangl2* knock-out**

**(A)** Schematic representation of the strategy to generate Emx1-Vangl2 mice. ‘Cre line’ (that expresses Cre recombinase under the control of Emx1 promoter) is crossed with a ‘Vangl2 flox/flox allele line’. Emx1-Vangl2 mice will express a truncated defective protein in the neurons and glia of the telencephalon starting at E10.5. **(B)** Western blots show a loss of expression of Vangl2 protein in the hippocampus of P21 Emx1-Vangl2 mice compared to the control. **(C)** High-magnification image of Vangl2 (red) and DCX (green) in the hippocampus shows that these proteins colocalise in the CA3 *stratum lucidum* (*sl*). High-magnification image of Vangl2 (red) and Tau (green) proteins colocalising in the *sl*. The high-magnification image shows that Vangl2 (red) and Znt3 (green) are present in the CA3 *sl* but do not colocalise. The high-magnification image of Vangl2 (red) and PSD-95 (green) labelling shows that these proteins do not colocalise in the *sl*. Scale bar, 25 µm.

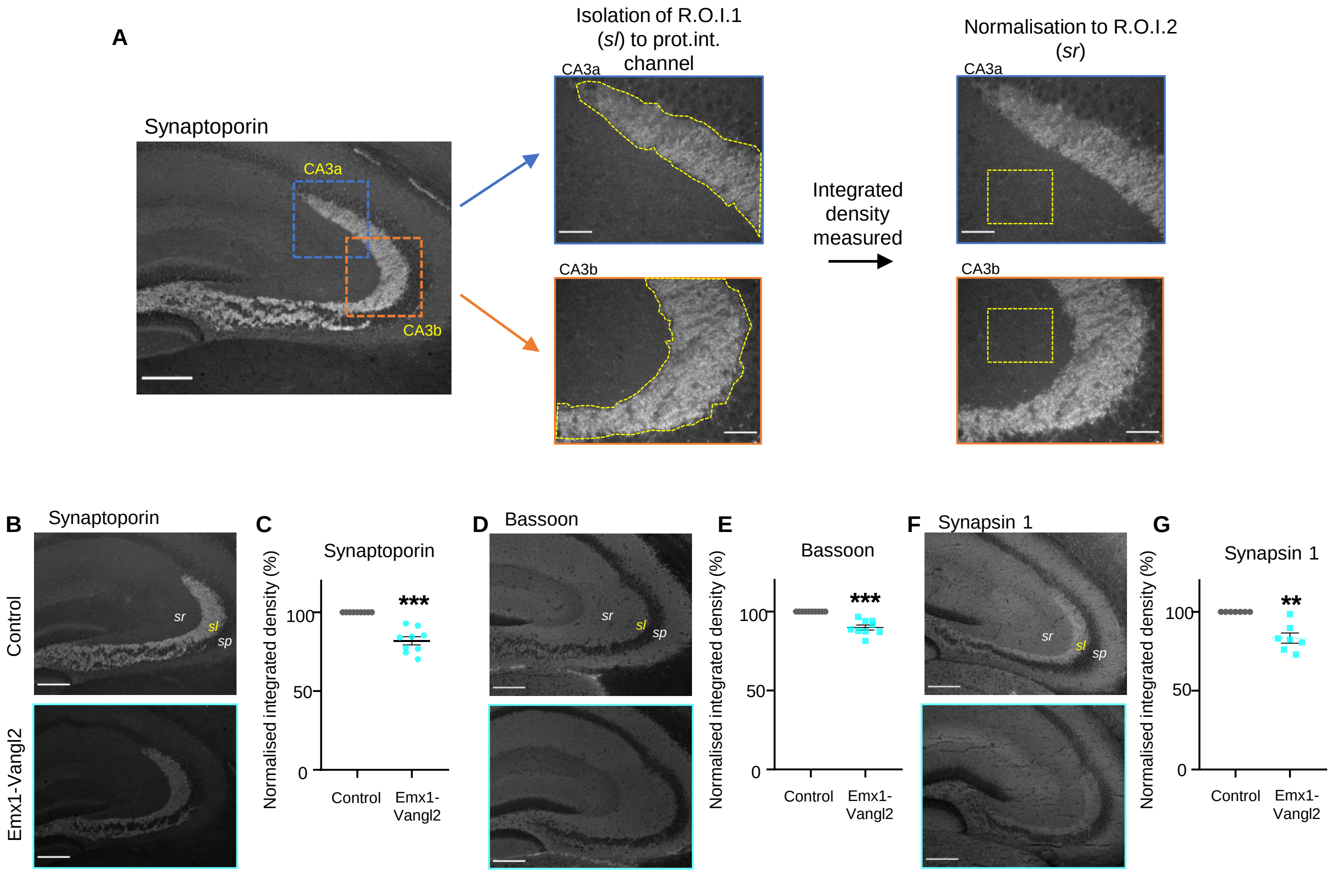


**Supplementary Fig. 2: Measure of integrated density of presynaptic markers.**

To quantify the intensity of the labellings, a mask corresponding to the *sl* was created in the red channel. This first region of interest (ROI) was applied to the green channel, where the integrated density of the protein of interest was measured only in the *sl*. The results are normalised onto values measured in a second ROI: the *stratum radiatum (sr)*. Scale bar, 200 µm. **(B, D, F)** Representative epifluorescence images of the *sl* with illustrations of labeling for synaptoporin (H), bassoon (J), and synapsin 1 (L) are depicted in P23 Emx1-Vangl2 and control mice. Scale bar, 200 µm. *sp: stratum pyramidale* **(C, E, G)** Quantification of the percentage of reduction of the normalised integrated density for the signal of synaptoporin (I), bassoon (K), and synapsin 1 (M) in P23 control (grey) and Emx1-Vangl2 (turquoise) mice. N control = 9 mice, N Emx1-Vangl2 = 9 mice. One sample t-test. $^{**}p < 0.01$, and $^{***}p < 0.001$. . From C to G data are presented as mean ±SEM and single data points are shown as dot.

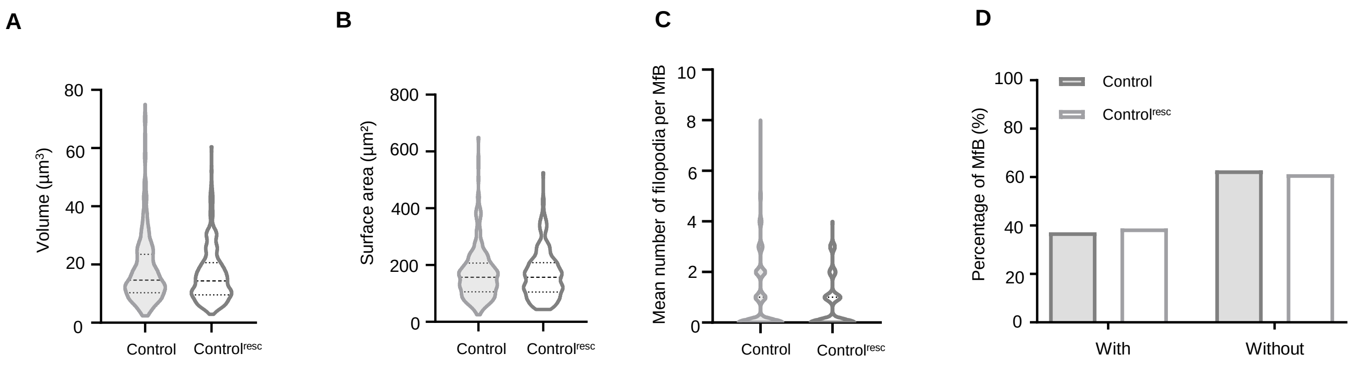


**Supplementary Fig. 3: Presynaptic re-expression of *vangl2* rescued the morphology of MfB in 10 week-old Emx1-Vangl2 mice.**
**(A-D)** Quantifications of volume (A), surface area (B) , number of filopodia per MfB (C) , and percentage of MfB with filopodia (D) in 10 week-old control (grey) and control$^{resc}$ (white) mice. N control = 271 MfB from 7 mice, N control$^{resc}$ = 263 MfB from 8 mice. Mann-Whitney test, ns p>0.05**.**

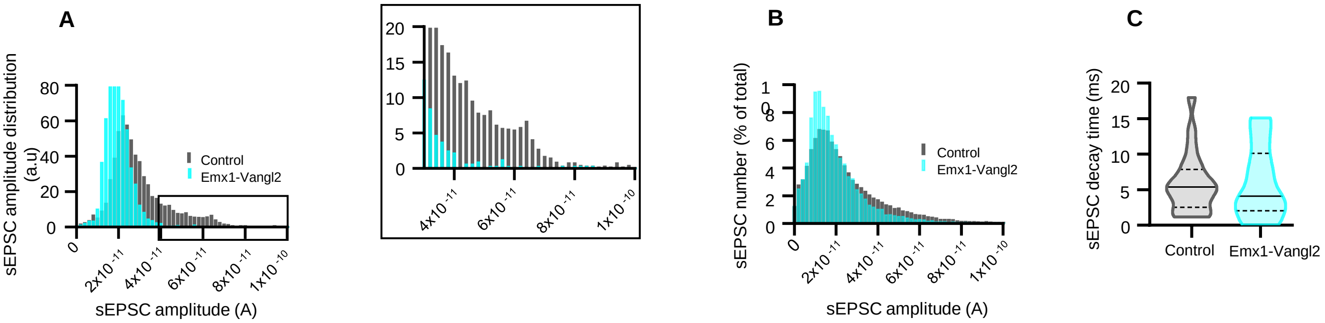


**Supplementary Fig. 4: Specific decrease of large sEPSC at MfB/TE synapses in Emx1-Vangl2 mice.**

**(A)** sEPSC amplitude distribution in CA3a pyramidal neurons in control (N = 16 mice, n = 23 cells) versus Emx1-Vangl2 (N = 14 mice, n = 19 cells). The binning size for the amplitude was 2 pA. The rectangle delimits the region where sEPSC above 30 pA is suppressed in Emx1-Vangl2 and is enlarged. **(B)** sEPSC number plotted against sEPSC amplitude. **(C)** sEPSC decay time (ms) in CA3a pyramidal neurons in P21-30 control (N =16 mice, n =23 cells) and Emx1-Vangl2 acute hippocampal slices (N= 14 mice, n = 19 cells). ns, p>0.05, Mann-Whitney test.

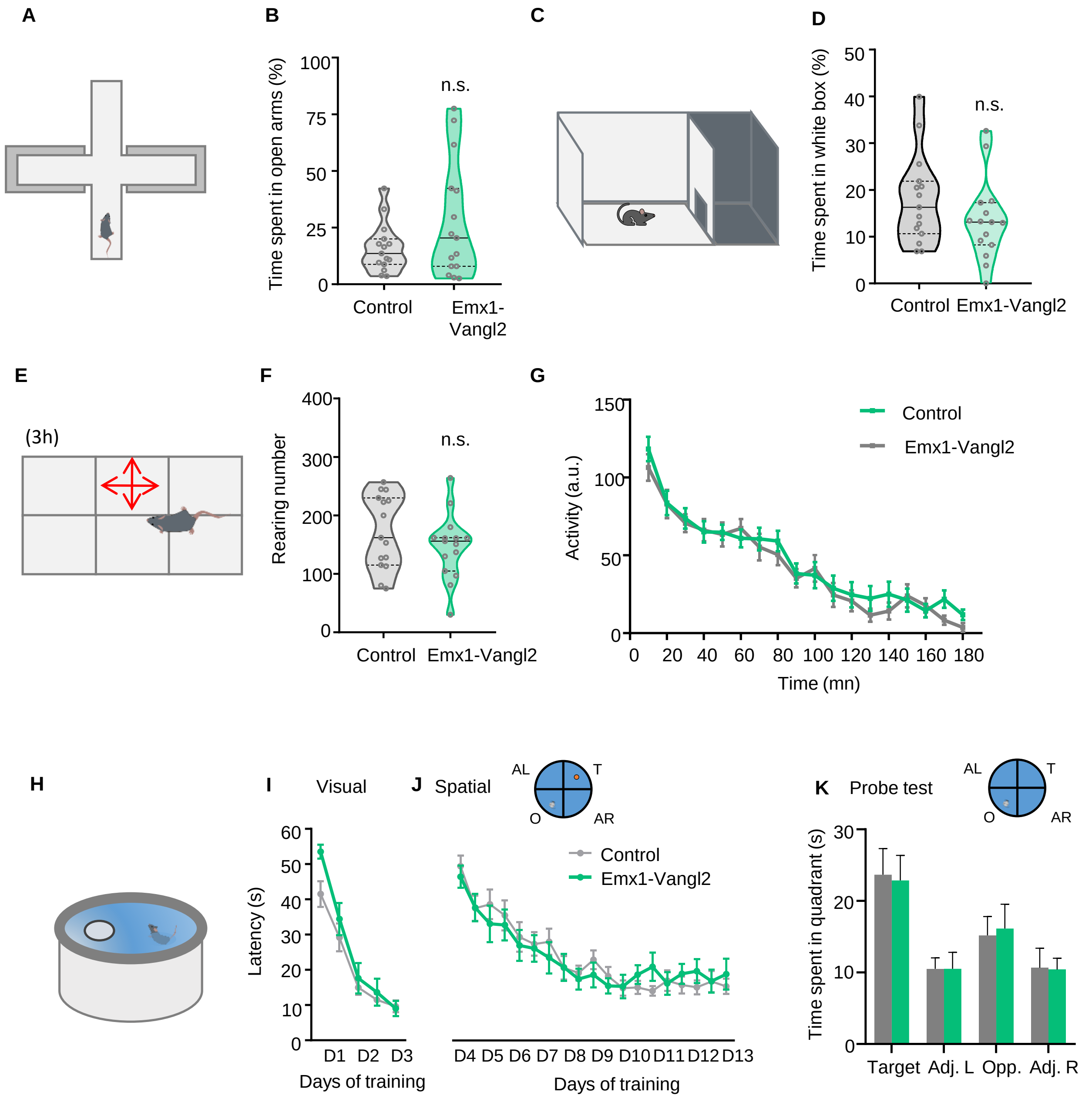


**Supplementary Fig. 5: Emx1-Vangl2 mice show neither anxiety-like behavior nor locomotor activity impairment as well as no spatial learning deficit.**

**(A)** Representation of the plus maze paradigm used to assess anxiety. **(B)** Quantification of the percentage of time spent in the open arms of the plus-maze test. Unpaired t-test, ns p>0.05. **(C)** Representation of the dark/light paradigm used to assess anxiety. **(D)** Quantification of the percentage of time spent in the white box of the dark/light test. Unpaired t-test, ns p>0.05. **(E)** Representation of the novel home cage paradigm used to assess spontaneous locomotor activity. **(F)** Quantification of the number of rearings during a 3h session in the novel home cage. Unpaired t-test, ns p>0.05. **(G)** Quantification of activity during a 3h session in the open field. Two-way ANOVA ns p>0.05. **(H)** Representation of the MWM paradigm used to assess spatial learning and memory. **(I-J)** Latency to escape the water maze test during the visual (I) and spatial (J) acquisition. Two-way ANOVA ns p>0.05. **(K)** Time spent in the four quadrants of the water maze during the probe test at 10 days during the spatial acquisition. T: Target, A.L: adjacent left, O: opposite, A.R: adjacent right. 2-way ANOVA, no genotype effect for any quadrants. Number of animals: control: n=15, Emx1-Vangl2: n=15.
